# Single-Organelle Quantification Reveals Stoichiometric and Structural Variability of Carboxysomes Dependent on the Environment

**DOI:** 10.1101/568238

**Authors:** Yaqi Sun, Adam J. M. Wollman, Fang Huang, Mark C. Leake, Lu-Ning Liu

## Abstract

The carboxysome is a complex, proteinaceous organelle that plays essential roles in carbon assimilation in cyanobacteria and chemoautotrophs. It comprises hundreds of protein homologs that self-assemble in space to form an icosahedral structure. Despite its significance in enhancing CO_2_ fixation and potentials in bioengineering applications, the formation of carboxysomes and their structural composition, stoichiometry and adaptation to cope with environmental changes remain unclear. Here we use live-cell single-molecule fluorescence microscopy, coupled with confocal and electron microscopy, to decipher the absolute protein stoichiometry and organizational variability of single β-carboxysomes in the model cyanobacterium *Synechococcus elongatus* PCC7942. We determine the physiological abundance of individual building blocks within the icosahedral carboxysome. We further find that the protein stoichiometry, diameter, localization and mobility patterns of carboxysomes in cells depend sensitively on the microenvironmental levels of CO_2_ and light intensity during cell growth, revealing cellular strategies of dynamic regulation. These findings, also applicable to other bacterial microcompartments and macromolecular self-assembling systems, advance our knowledge of the principles that mediate carboxysome formation and structural modulation. It will empower rational design and construction of entire functional metabolic factories in heterologous organisms, for example crop plants, to boost photosynthesis and agricultural productivity.

**One Sentence Summary:** Determination of absolute protein stoichiometry reveals the organizational variability of carboxysomes in response to microenvironmental changes

The authors responsible for distribution of materials integral to the findings presented in this article in accordance with the policy described in the Instructions for Author (www.plantcell.org) is: Lu-Ning Liu.

## INTRODUCTION

Organelle formation and compartmentalization within eukaryotic and prokaryotic cells provide the structural foundation for segmentation and modulation of metabolic reactions in space and time. Bacterial microcompartments (BMCs) are self-assembling organelles widespread among bacterial phyla (Axen et al., 2014). By physically sequestering specific enzymes key for metabolic processes from the cytosol, these organelles play important roles in CO_2_ fixation, pathogenesis, and microbial ecology (Yeates et al., 2010; Bobik et al., 2015). According to their physiological roles, three types of BMCs have been characterized: the carboxysomes for CO_2_ fixation, the PDU microcompartments for 1,2−propanediol utilization, and the EUT microcompartments for ethanolamine utilization.

The common features of various BMCs are that they are ensembles composed of purely protein constituents and comprise an icosahedral single-layer shell that encases the catalytic enzyme core. This proteinaceous shell, structurally resembling virus capsids, is self-assembled from several thousand polypeptides of multiple protein paralogs that form hexagons, pentagons and trimers (Kerfeld and Erbilgin, 2015; Sutter et al., 2016; Faulkner et al., 2017). The highly-ordered shell architecture functions as a physical barrier that concentrates and protects enzymes, as well as selectively gating the passage of substrates and products of enzymatic reactions (Yeates et al., 2010; Bobik et al., 2015).

Carboxysomes serve as the key CO_2_-fixing machinery in all cyanobacteria and some chemoautotrophs. The primary carboxylating enzymes, ribulose-1,5-bisphosphate carboxylase oxygenase (Rubisco) (Rae et al., 2013), are encapsulated by the carboxysome shell that facilitates the diffusion of HCO_3_^−^ and probably reduces CO_2_ leakage into the cytosol (Dou et al., 2008). Based on the form of enclosed Rubisco, carboxysomes can be categorized into two different classes, α-carboxysomes and β-carboxysomes (Rae et al., 2013; Kerfeld and Melnicki, 2016). The β-carboxysomes in the cyanobacterium *Synechococcus elongatus* PCC7942 (Syn7942) have been extensively characterized as the model carboxysomes. The shell of β-carboxysomes from Syn7942 is composed of the hexameric proteins CcmK2, CcmK3 and CcmK4 that form predominately the shell facets (Kerfeld et al., 2005), the pentameric protein CcmL that caps the vertices of the polyhedron (Tanaka et al., 2008), as well as the trimeric proteins CcmO and CcmP (Cai et al., 2013; Larsson et al., 2017). The core enzymes of β-carboxysomes consist of a paracrystalline arrangement of plant-type

Rubisco (comprising the large and small subunits RbcL and RbcS) and β-carbonic anhydrase (β-CA, encoded by the *ccaA* gene). The colocalized β-CA dehydrates HCO_3_^−^ to CO_2_ and creates a CO_2_-rich environment in the carboxysome lumen to favor the carboxylation of Rubisco. In addition, CcmM and CcmN function as “linker” proteins to promote Rubisco packing and shell-interior association (Kinney et al., 2012). CcmM in the β-carboxysome appears as two isoforms, a 35-kDa truncated CcmM35 and a full-length 58-kDa CcmM58 (Long et al., 2007; Long et al., 2010; Long et al., 2011). CcmM35 contains three Rubisco small subunit-like (SSU) domains that interact with Rubisco (Hagen et al., 2018b; Wang et al., 2019), whereas CcmM58 has an N-terminal γ-CA-like domain in addition to the SSU domains and recruits CcaA to the shell. RbcX is recognized as a chaperonin-like protein for Rubisco assembly (Emlyn-Jones et al., 2006; Saschenbrecker et al., 2007; Occhialini et al., 2016); it has been recently revealed to serve as one component of the carboxysome and play roles in mediating carboxysome assembly and subcellular distribution (Huang et al., 2019).

Understanding the physiological composition and assembly principles of carboxysome building blocks is of key importance not solely to unravel the underlying molecular mechanisms of carboxysome formation and biological functions, but also for heterologously engineering and modulating functional CO_2_-fixing organelles to supercharge photosynthetic carbon fixation in synthetic biology applications. Previous estimations of the carboxysome protein stoichiometry from either the whole cell lysates or the isolated forms using immunoblot and mass spectrometry illustrated the relative abundance of carboxysome proteins (Long et al., 2005; Long et al., 2011; Rae et al., 2012; Faulkner et al., 2017). Moreover, it was revealed that carboxysome biosynthesis in Syn7942 is highly dependent upon environmental conditions during cell growth, such as light intensity (Sun et al., 2016) and CO_2_ availability (McKay et al., 1993; Harano et al., 2003; Woodger et al., 2003; Whitehead et al., 2014). The exact stoichiometry of all building components in the functional carboxysome and how carboxysomes manipulate their compositions, organizations and functions to cope with environmental changes have remained elusive.

Here, we construct a series of Syn7942 mutants with individual components of carboxysomes functionally tagged with the bright and fast-maturing enhanced yellow fluorescent protein (YFP) and report the *in vivo* characterization of protein stoichiometry of carboxysomes at the single-organelle level, using real-time single-molecule fluorescence microscopy, confocal and electron microscopy combined with a suite of biochemical and genetic assays. Quantification of the protein stoichiometry of β-carboxysomes in Syn7942 grown under different conditions demonstrates the organizational flexibility of β-carboxysomes, and their ability to modulate functions towards local alterations of CO_2_ levels and light intensity during cell growth, as well as the regulation of the spatial localization and mobility of β-carboxysomes in the cell. This study provides fundamental insight into the formation and structural plasticity of carboxysomes and their dynamic organization towards environmental changes, which could be extended to other BMCs and macromolecular systems. A deeper understanding of the protein composition and structure of carboxysomes will inform strategies for rational design and engineering of functional and adjustable metabolic modules towards biotechnological applications.

## RESULTS

### Protein stoichiometry of functional carboxysomes at the single-organelle level

We constructed ten Syn7942 strains expressing individual β-carboxysome proteins (CcmK3, CcmK4, CcmK2, CcmL, CcmM, CcmN, RbcL, RbcS, CcaA, RbcX) fused with YFP at their C-termini individually (Supplemental Figure 1). Fluorescence tagging at the native chromosomal locus under the control of their native promoters ensures expression of the fluorescently-tagged proteins in context and at physiological levels (Sun et al., 2016). Eight of these strains, in which YFP was fused to CcmK3, CcmK4, CcmL, CcmM, CcmN, RbcS, CcaA, and RbcX respectively, are fully segregated (Supplemental Figures 1C and 2) and exhibit wild-type levels of cell size, growth and carbon fixation within experimental error (Supplemental Table 1), consistent with previous observations (Savage et al., 2010; Cameron et al., 2013; Sun et al., 2016; Faulkner et al., 2017; Huang et al., 2019).

**Table 1.** Protein stoichiometry of the Syn7942 β-carboxysome and its variability in cells grown under Air/ML, CO_2_/ML, LL and HL conditions determined from Slimfield and confocal microscopy. Stoichiometry is presented as Peak value ± HWHM and the sample sizes are indicated as *n*. Peak values were determined from Slimfield stoichiometry profiles of each carboxysome proteins (Figure 1, Supplemental Figure 3). Quantification of CcmL under the four conditions was acquired from Slimfield for accurate measurement of copies of shell pentamers for capping the carboxysome structure. Copies of other carboxysome proteins were calculated using Slimfield results (bold) with definitive counts of protein copies under Air/ML (See also Supplemental Figure 3) in combination with relative quantification of each protein under the four conditions from confocal imaging (See also Supplemental Figure 7 and 8). Protein structures were derived from previous studies (Kerfeld et al., 2005; Long et al., 2007; Tanaka et al., 2007; Tanaka et al., 2008; Long et al., 2011; Kinney et al., 2012; McGurn et al., 2016). *Monomeric unit of CcmM was designated to CcmM35 that is the majority of CcmM; CcmM58 is postulated as a trimer.

| Category | Structure | Protein | Air/ML |  | CO<sub>2</sub>/ML |  | LL |  | HL |  |
| --- | --- | --- | --- | --- | --- | --- | --- | --- | --- | --- |
| | | | Peak value ± HWHM | Number of functional units | Peak value ± HWHM | Number of functional units | Peak value ± HWHM | Number of functional units | Peak value ± HWHM | Number of functional units |
| Shell proteins | Hexamer | CcmK3 | **92 ± 148**<br>(*n* = 219) | **15 ± 25** | 172 ± 83<br>(*n* = 2048) | 29 ± 14 | 83 ± 31<br>(*n* = 1516) | 14 ± 5 | 87 ± 52<br>(*n* = 2155) | 14 ± 9 |
| | | CcmK4 | **314 ± 194**<br>(*n* = 77) | **52 ± 32** | 562 ± 263<br>(*n* = 1918) | 94 ± 44 | 313 ± 121<br>(*n* = 1766) | 52 ± 20 | 304 ± 95<br>(*n* = 3215) | 51 ± 16 |
|  | Pentamer | CcmL | **37 ± 17**<br>(*n* = 316) | **7.4 ± 3.4** | **66 ± 24**<br>(*n* = 311) | **13.2 ± 4.8** | **34 ± 15**<br>(*n* = 394) | **6.8 ± 3.0** | **69 ± 24**<br>(*n* = 220) | **13.8 ± 4.8** |
| Structural proteins | Monomer* | CcmM | **719 ± 1433**<br>(*n* = 71) | **719 ± 1433** | 468 ± 425<br>(*n* = 2313) | 468 ± 425 | 483 ± 366<br>(*n* = 3655) | 483 ± 366 | 1176 ± 691<br>(*n* = 2318) | 1176 ± 691 |
| | Monomer | CcmN | **74 ± 51**<br>(*n* = 86) | **74 ± 51** | 52 ± 28<br>(*n* = 3143) | 52 ± 28 | 51 ± 20<br>(*n* = 4022) | 51 ± 20 | 82 ± 34<br>(*n* = 5074) | 82 ± 34 |
| CA | Hexamer | CcaA | **86 ± 81**<br>(*n* = 95) | **14 ± 14** | 129 ± 86<br>(*n* = 1354) | 21 ± 14 | 65 ± 21<br>(*n* = 217) | 11 ± 4 | 122 ± 59<br>(*n* = 2837) | 20 ± 10 |
| Rubisco enzyme | L<sub>8</sub>S<sub>8</sub> | RbcS | **6822 ± 9200**<br>(*n* = 60) | **853 ± 1150** | 4401 ± 6655<br>(*n* = 894) | 550 ± 832 | 2934 ± 5492<br>(*n* = 752) | 367 ± 687 | 12057 ± 5186<br>(*n* = 1974) | 1507 ± 648 |
| Rubisco chaperone | Dimer | RbcX | **39 ± 32**<br>(*n* = 211) | **20 ± 16** | 38 ± 10<br>(*n* = 1370) | 19 ± 5 | 40 ± 9<br>(*n* = 1402) | 20 ± 5 | 40 ± 9<br>(*n* = 1861) | 20 ± 5 |

By contrast, RbcL-YFP and CcmK2-YFP strains are only partially segregated, in agreement with previous studies (Savage et al., 2010; Cameron et al., 2013; Sun et al., 2016). Through immunoblot analysis using anti-fluorescence protein, anti-RbcL and anti-CcmK2 antibodies (Supplemental Figure 2B), we estimate that 29.2 ± 7.1 % (mean ± standard deviation (SD), *n* = 4) of total RbcL and 6.0 ± 0.7 % (*n* = 3) of total CcmK2 were tagged with YFP in RbcL-YFP and CcmK2-YFP strains. Nevertheless, we excluded the stoichiometric quantification of RbcL and CcmK2 in this study, in view of the partial segregation which could result in quantification inaccuracy.

We used single-molecule Slimfield microscopy (Plank et al., 2009) to visualize individual carboxysomes that were fused with YFP (Figure 1, Supplemental Figure 3). This technique allows detection of fluorescently-labelled proteins with millisecond sampling, enabling real-time tracking of rapid protein dynamics inside living cells, exploited previously to study functional proteins involved in bacterial DNA replication and remodeling (Reyes-Lamothe et al., 2010; Badrinarayanan et al., 2012), gene regulation in budding yeast cells (Wollman et al., 2017; Leake, 2018), bacterial cell division (Lund et al., 2018), and chemokine signaling in lymph nodes (Miller et al., 2018). Our prior measurements using relatively fast-maturing fluorescent proteins such as YFP suggest that less than 15% of fluorescent proteins are likely to be in a non-fluorescent immature state (Leake et al., 2008; Shashkova et al., 2018).

**Figure 1.**
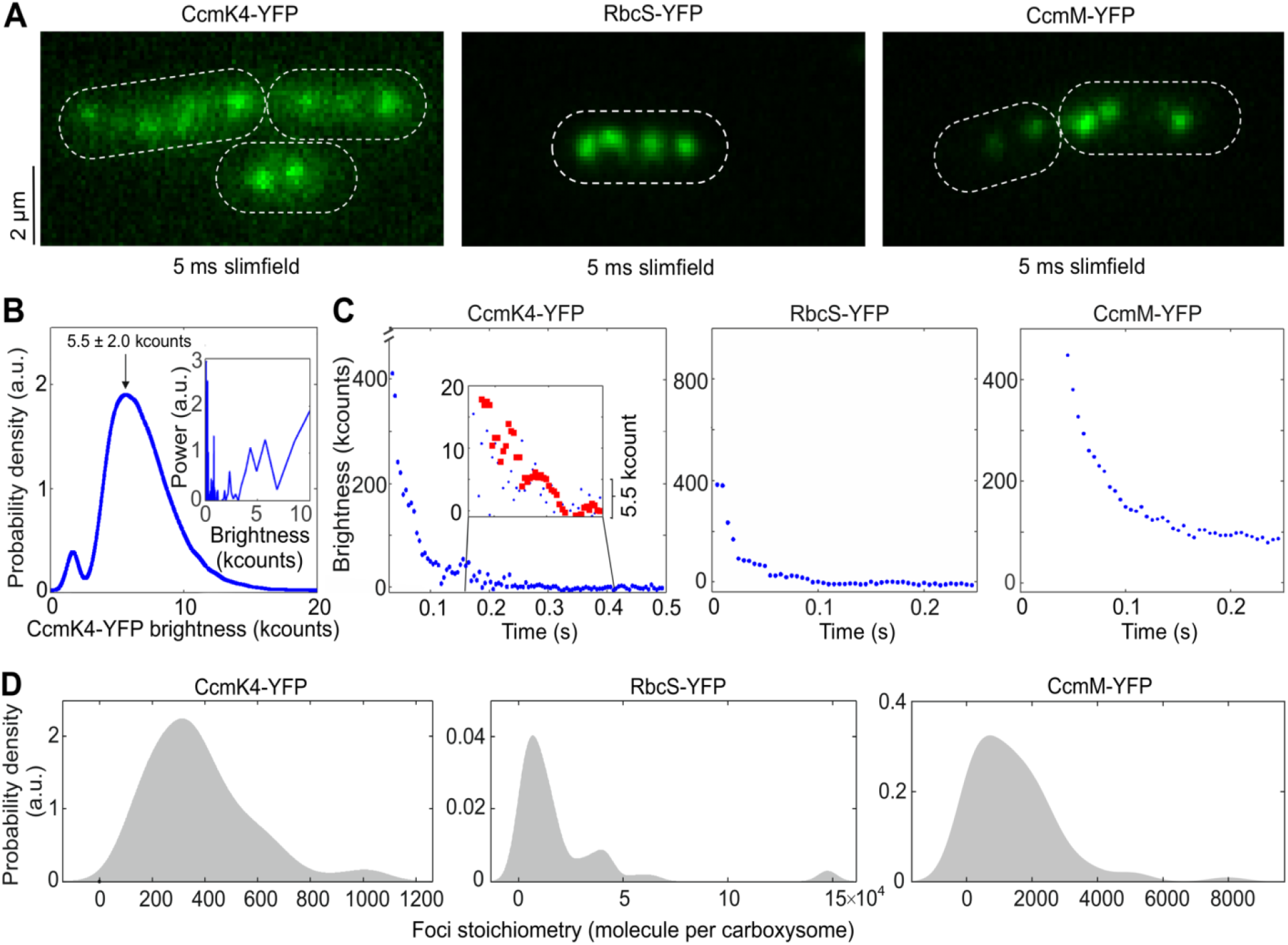
Slimfield quantification of cells grown under ambient air/moderate light Air/ML conditions. **(A)** Averaged Slimfield images of YFP fluorescence (green) over 5 frames of strains expressing shell component CcmK4-YFP, the interior enzyme RbcS-YFP, and the shell-interior linker protein CcmM-YFP. White dashed lines indicate cell body outlines. **(B)** Distribution of the intensities of automatically detected foci from the end of photobleaching, corresponding to the characteristic intensity of *in vivo* YFP. Inset shows the Fourier spectrum of ‘overtracked’ foci, tracked beyond photobleaching, showing a peak at the characteristic intensity. **(C)** Representative fluorescence photobleaching tracked at ultra-fast speed. The CcmK4 plot shows an inset ‘zoomed in’ on lower intensity range with step-preserving Chung-Kennedy filtered data in red, showing individual photobleaching steps clearly visible at the characteristic intensity. Brightness (kcounts), counts measured per camera pixel multiplied by 1,000. **(D)** Distribution of YFP copy number detected for individual carboxysomes in corresponding mutants, rendered as kernel density estimates using standard kernel width. Heterogeneity of contents was observed, and a “preferable” copy number, represented by kernel density peak values could be determined. Statistics of copy numbers (Peak value ± HWHM) are listed in Table 1 for ML conditions. The corresponding Slimfield images and histogram for complete strain sets are shown in Supplemental Figure 3.

Figure 1A shows the Slimfield images of three representative Syn7942 strains RbcS-YFP, CcmK4-YFP and CcmM-YFP that grow under ambient air and moderate light (hereafter denoted Air/ML), to determine the protein stoichiometry from different carboxysome structural domains. Single carboxysomes are detected as distinct fluorescent foci in cells of the YFP-fused strains (Figure 1A, Supplemental Figure 3), whose sigma width is approximately 250 nm (*n* = 100), comparable to the diffraction-limited point spread function width of our imaging system. We use the number of YFP molecules per fluorescent focus as an indicator of the stoichiometry of the fluorescently-labelled protein subunits in each individual carboxysomes, which we determined by quantifying step-wise photobleaching of the fluorescent tag during the Slimfield laser excitation process (Figures 1B to 1C, Table 1) using a combination of Fourier spectral analysis and edge-preserving filtration of the raw data (Leake et al., 2003; Leake et al., 2004; Leake et al., 2006) (see details in Materials and Methods). The resulting broad distributions of protein stoichiometry, rendered as kernel density estimates, suggest a variable content of individual components per carboxysome (Figure 1D), indicative of the structural heterogeneity of β-carboxysomes. The modal average stoichiometry of each protein subunit per carboxysome was defined by the measured peak from each distribution of the raw stoichiometric data (Figure 1D, Supplemental Figure 3), after subtracting the background fluorescence distribution, primarily from chlorophylls, which was determined from the WT cells (Supplemental Figure 4).

In the β-carboxysome synthesized in cells grown under Air/ML, Rubisco enzymes are the predominant components, as indicated by the RbcS content (Table 1). CcmM is the second most abundant element; there are over 700 copies of CcmM molecules per β-carboxysome. In addition, the CcmK4 content is greater than that of CcmK3 by a factor of 3.8. CcmL, CcmN, CcaA and RbcX are the minor components in the β-carboxysome. Our results reveal that there are 37 CcmL subunits per carboxysome, with the raw stoichiometry distribution showing some indications of peaks at multiples of ∼5 molecules indicative of multiples of CcmL pentamers (Supplemental Figure 4C), consistent with the atomic structure of CcmL (Tanaka et al., 2008). A modal average of 37 CcmL molecules thus suggests that a single carboxysome contains an average of 7.4 CcmL pentamers, less than the 12 CcmL pentamers that were postulated to occupy all the vertices of the icosahedral shell (Bobik et al., 2015; Kerfeld et al., 2018). It is feasible that not all vertices of the carboxysome structure are capped by CcmL pentamers, as BMC shells deficient in pentamers could still be formed without notable structural variations (Cai et al., 2009; Lassila et al., 2014; Hagen et al., 2018a). Our study represents a direct characterization of protein stoichiometry at the level of single functional carboxysomes in their native cellular environment.

As a control, we fused RbcL with mYPet, a monomeric-optimized variant of YFP. The RbcL-YFP and RbcL-mYPet cells show no significant difference in the subcellular distribution of carboxysomes as well as cell doubling times and carbon fixation (Supplemental Figure 5), demonstrating that there are no measurable artefacts due to putative effects of dimerization of the YFP tag.

We also examined the relative abundance of individual carboxysome proteins in the YFP-fusion Syn7942 strains in cell lysates, using immunoblot probing with an anti-fluorescent protein antibody (Supplemental Figure 2A, Supplemental Table 2). To compare with the stoichiometry obtained from Slimfield, we normalized the abundance of carboxysome proteins estimated from immunoblot analysis, using the RbcS content per carboxysome determined by Slimfield. It appears that the content of β-carboxysome proteins determined by immunoblotting is generally greater than that within the carboxysome characterized by Slimfield. Despite the potential effects caused by YFP fusion, this could suggest the presence of a “storage pool” of carboxysome proteins located in the cytoplasm that are involved in the biogenesis, maturation and turnover of carboxysomes. The ratio of RbcL/S detected from cell lysates fraction is about 8:5.8 (*n* = 4) (Supplemental Table 2), in line with previous results (Long et al., 2011) but distinct from the *in vitro* reconstitution observations (Ryan et al., 2018; Wang et al., 2019).

### Stoichiometry of carboxysome proteins exhibit a dependence on the microenvironment conditions of live cells

Our previous study showed that the content and spatial positioning of β-carboxysomes in Syn7942 are dependent upon light intensity during cell growth, revealing the physiological regulation of carboxysome biosynthesis (Sun et al., 2016). Whether the stoichiometry of different components in the carboxysome structure changes in response to fluctuations in environmental conditions is unknown. Here we addressed this question by taking advantage of the far greater throughput of confocal microscopy compared to Slimfield, whilst still using the single-molecule precise Slimfield data as a calibration to convert the intensity of detected foci from confocal images into estimates for absolute numbers of stoichiometry. We achieved this by identifying the peak value of the foci intensity distribution from each given cell strain obtained from confocal imaging with the peak value of the measured Slimfield foci stoichiometry distribution for the equivalent cell strain under Air/ML. This approach allows us to generate a conversion factor which we then applied to subsequent confocal data acquired under lower light (LL), higher light (HL) and ML with the air supplemented by 3% CO_2_, and to estimate relative changes in the stoichiometry of carboxysome building components using large numbers of cells, without the need to obtain separate Slimfield datasets for each condition (Figure 2, Supplemental Figures 6 to 8).

**Figure 2.**
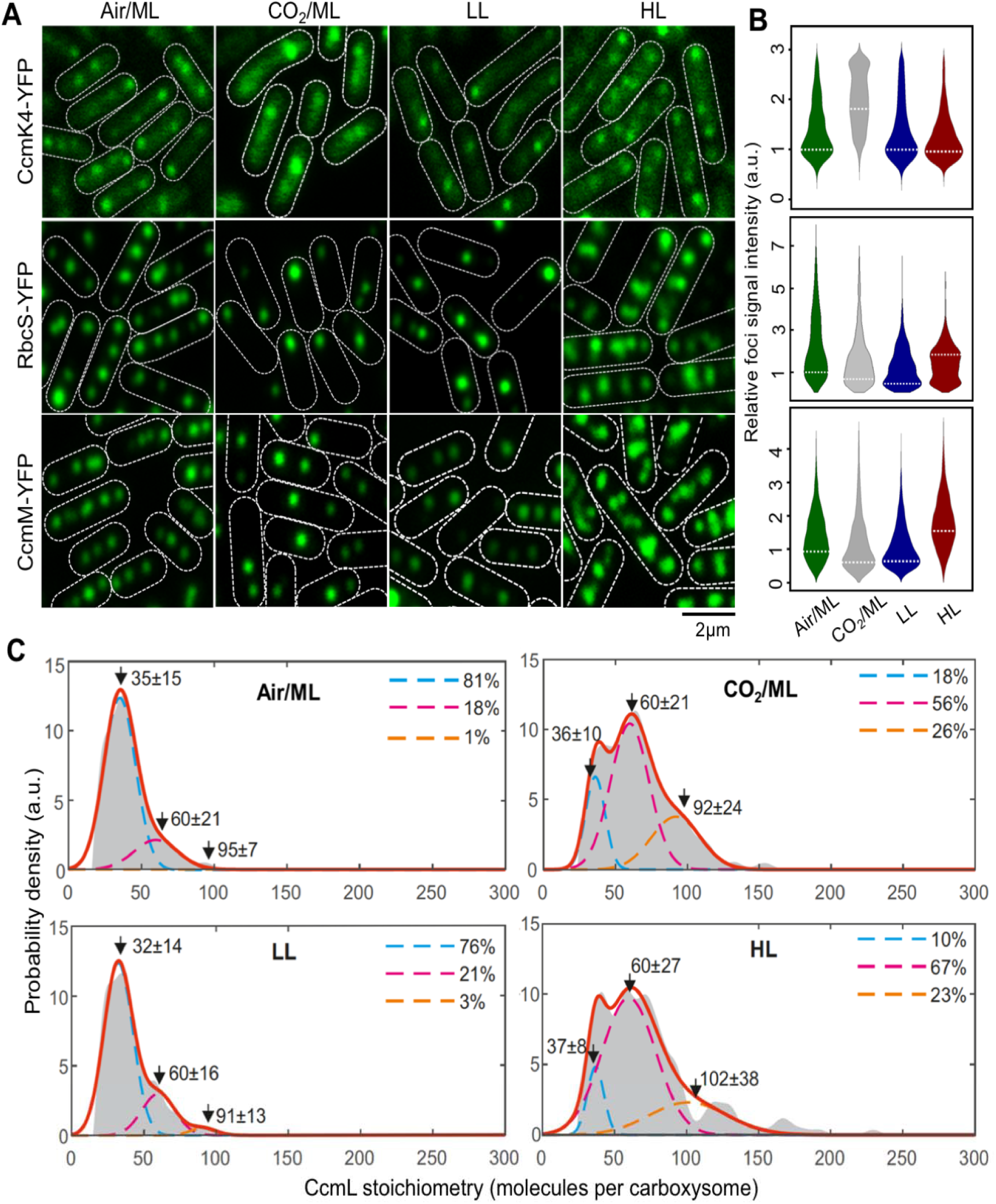
Relative protein quantification of CcmK4, RbcS and CcmM in the carboxysome under different CO_2_ levels and light intensities using confocal microscopy. **(A)** Confocal images of CcmK4-YFP, RbcS-YFP and CcmM-YFP strains under Air/ML, CO_2_/ML, LL and HL. Fluorescence foci (green) indicate carboxysomes, and cell borders were outlined by white dashed lines. Scale bar indicates 2 μm. **(B)** Violin plot of carboxysome intensities under Air/ML, CO_2_/ML, LL and HL, normalized to kernel density ML peak values (peaks marked by white dashed lines). **(C)** Kernel density estimates of CcmL carboxysome copy number grown under Air/ML, CO_2_, LL and HL detected by Slimfield and corrected for chlorophyll. Triple Gaussian fits are indicated as colored dashed lines with the summed fit in red. The percentage in each Gaussian is indicated aside.

Figure 2A shows confocal fluorescence images of RbcS-YFP, CcmK4-YFP, and CcmM-YFP strains grown under Air/ML, 3% CO_2_ (CO_2_/ML), LL and HL. The confocal images reveal classic patterns of cellular localization of carboxysomes similar to those observed with Slimfield microscopy (Supplemental Figure 6). We analyzed the confocal images to detect carboxysome fluorescent foci within the cells and quantify their fluorescence intensities (Figure 2B, Supplemental Figures 7 and 8). We find that the number of carboxysomes per cell is dependent on growth conditions: it is reduced under CO_2_/ML in contrast to Air/ML, whereas HL increases the abundance of β-carboxysomes (Supplemental Table 3), consistent with previous findings (Whitehead et al., 2014; Sun et al., 2016). The slightly different carboxysome contents estimated in individual YFP-fused strains might suggest potential mechanisms of the cells that tune carboxysome organization. As a common feature, the abundance of all the proteins in the β-carboxysome is apparently modulated under distinct growth conditions. For instance, both RbcS and CcmM have a higher content per carboxysome under HL compared with that under other conditions, whereas the CcmK4 content per β-carboxysome increase under 3% CO_2_ (Figure 2B). The dependence of carboxysome protein stoichiometry inferred from the peak values of the stoichiometry distributions under different cellular microenvironmental conditions is summarized in Table 1.

Interestingly, we find that the variation of CcmL abundance per carboxysome rises with increasing light illumination and CO_2_ availability (Figure 2C). The measured stoichiometry distribution of CcmL pentamers suggests the presence of three populations: (I) carboxysomes with < 60 CcmL subunits (in the range of 32-37); (II) carboxysomes with 60 CcmL subunits, consistent with the expectation that 12 vertices of the icosahedral carboxysome are fully occupied by CcmL pentamers (Tanaka et al., 2008; Rae et al., 2013; Kerfeld et al., 2018); (III) carboxysomes with > 60 CcmL subunits (in the range of 91-102). Using a nearest-neighbor model to estimate the probability for the diffraction-limited optical images of individual carboxysomes in a cell, we find that the Population III carboxysomes represent random overlap of two or more carboxysomes from the Population I and II (Figure 2C). Population I represents a “non-complete capped” state in which not all vertices in the icosahedron are occupied by CcmL pentamers. We find the characteristic stoichiometry of the Population I carboxysomes increases with the enhancement of light intensity during cell growth, from 32 CcmL molecules (LL) to 35 (ML) and 37 (HL), with HL having a significantly smaller proportion (23%) of “non-complete capped” carboxysomes compared to ∼80% under LL and ML conditions. Supplementing the air with 3% CO_2_ under ML similarly results in a substantial decrease in the proportion of “non-complete capped” carboxysomes in the population (18%) comparable to the HL condition in the absence of any supplemental CO_2_. These findings suggest a dependence of carboxysome assembly which may allow adaptation towards microenvironmental changes, i.e. the increase in the population of capped carboxysomes in situations which are favorable towards photosynthesis (HL conditions and locally-raised levels of CO_2_).

This finding is also validated by the changes in protein abundance of other carboxysome components under environmental regulation (Table 1, Supplemental Figures 7 and 8). Cells were maintained under different growth conditions prior to microscopy imaging, to ensure their full acclimation. Variations of protein content in carboxysomes under CO_2_/ML *vs*. Air/ML, and HL *vs*. LL conditions indicate distinct fashions of stoichiometric regulation of carboxysome building blocks (Figure 3, Supplemental Table 4). The abundance of CcmK3 and CcmK4, whose encoding genes are distant from the *ccmKLMNO* operon (Sommer et al., 2017), increases under 3% CO_2_ and remains relatively constant under HL/LL, contrary to the changes in the abundance of CcmN and CcmM that are located in the *ccm* operon. In addition, the ratio of CcmK4:CcmK3 per carboxysome appear to be relatively constant, in the range of 3.6−4.1 (Supplemental Table 5), indicating the organizational correlation between CcmK3 and CcmK4 within the β-carboxysome structure. We find the rise of CcaA content and reduction of RbcS content under CO_2_/ML *vs*. Air/ML, whereas both increase under HL, suggesting distinct regulation of the two components. It has been recently demonstrated that the putative Rubisco chaperone RbcX is part of the carboxysome and plays roles in mediating carboxysome formation (Huang et al., 2019). The fold changes of RbcX content in each carboxysome under different conditions are close to 1 (Figure 3), probably ascribed to the fact that its encoding gene is distant from the *rubisco* and *ccm* operons in Syn7942. Collectively, these results highlight the highly flexible stoichiometry of individual components within the natural carboxysomes in response to environmental changes.

**Figure 3.**
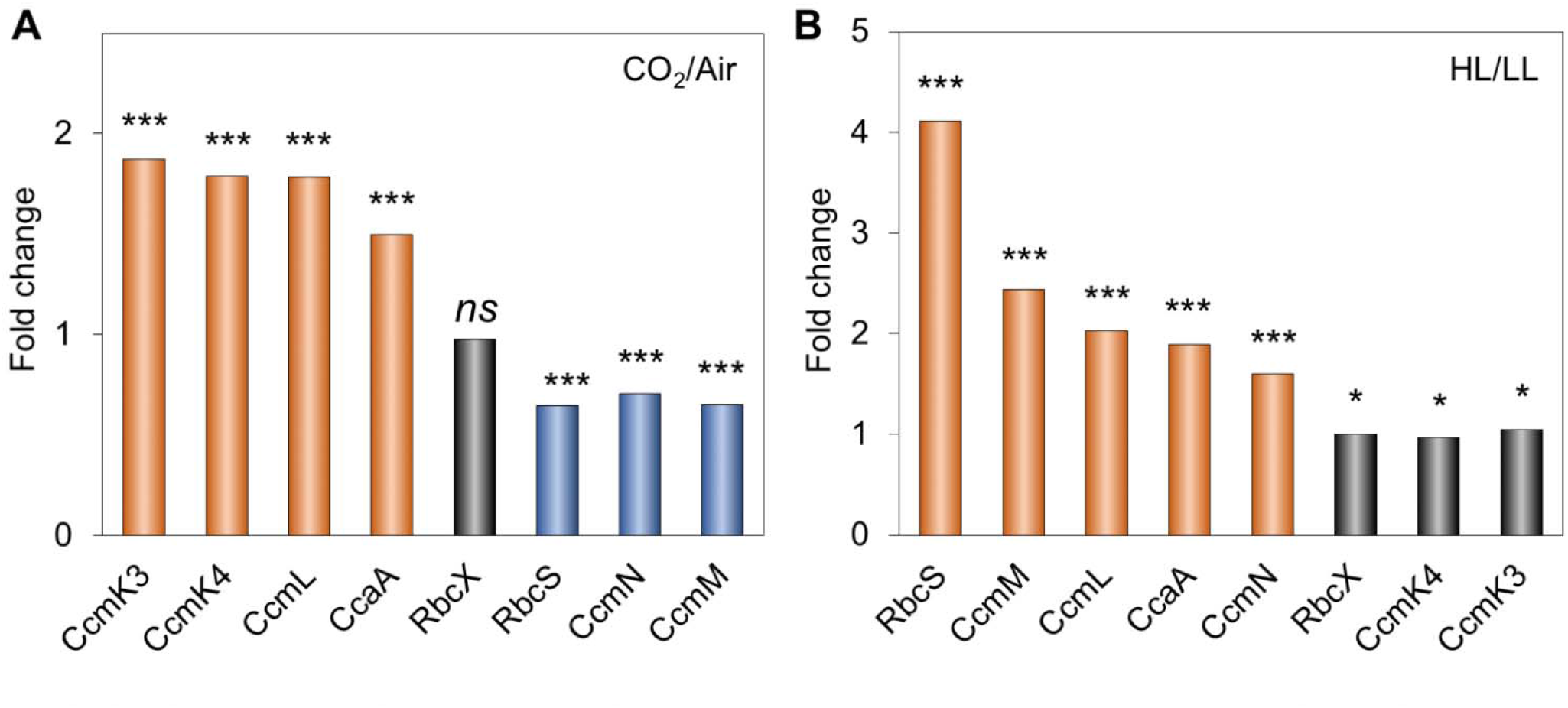
Changes in carboxysome protein stoichiometry upon increases in CO_2_ levels and light intensity. **(A)** Comparison of carboxysome protein stoichiometry under CO_2_ treatment. Increase in the CO_2_ concentration resulted in the rise of CcmK3, CcmK4, CcaA and CcmL contents and the decline of RbcS, CcmN and CcmM contents. **(B)** Comparison of carboxysome protein stoichiometry under light intensity treatment. Increased light intensity led to the elevation of RbcS, CcmM, CcmL, CcaA and CcmN contents, whereas the abundance of RbcX, CcmK3 and CcmK4 contents per carboxysome did not change dramatically. Mann-Whitney U-tests were performed to compare the numbers of functional units of individual carboxysome proteins changed from CO_2_/ML to Air/ML (A) and from HL to LL (B). *, *p* < 0.05; ***, *p* < 0.005; *ns, p* > 0.05.

### Variation of carboxysome diameter represents a strategy for manipulating carboxysome activity to adapt to environmental conditions

The change in the protein content per carboxysome signifies the variation of β-carboxysome size and organization among different cell growth conditions. Indeed, electron microscopy (EM) of Syn7942 WT cells substantiates the variable structures of β-carboxysomes in response to the changing environment (Figures 4A and 4B). The average diameter of β-carboxysomes is 192 ± 41 nm (*n* = 33) in Air/ML, 144 ± 24 nm (*n* = 25) in 3% CO_2_, 151 ± 22 nm (*n* = 27) in LL, and 208 ± 28 nm (*n* = 51) in HL (Figure 4B, Supplemental Table 5, Supplemental Figure 9). These results reveal that both the CO_2_ level and light intensity can result in alternations of carboxysome size (Figure 4B). Larger β-carboxysomes can accommodate more Rubisco enzymes (estimated on the basis of RbcS content) (Figure 4C). An exception is the carboxysomes under LL, which are around 5% larger than the carboxysomes under 3% CO_2_ but comprises only 67% of Rubisco per carboxysome under CO_2_ (Figure 4C, Supplemental Table 5). EM images reveal that the lumen of β-carboxysomes synthesized under LL often contain regions with low protein density (Figure 4A, arrows; Supplemental Figure 9), 59% for LL (16 out of 27 carboxysomes) compared with 9% for Air/ML (3 out of 33), 12% for CO_2_/ML (3 out of 25) and 8% for HL (4 out of 51), which likely accounts for the reduced and uneven Rubisco loading within the β-carboxysome.

**Figure 4.**
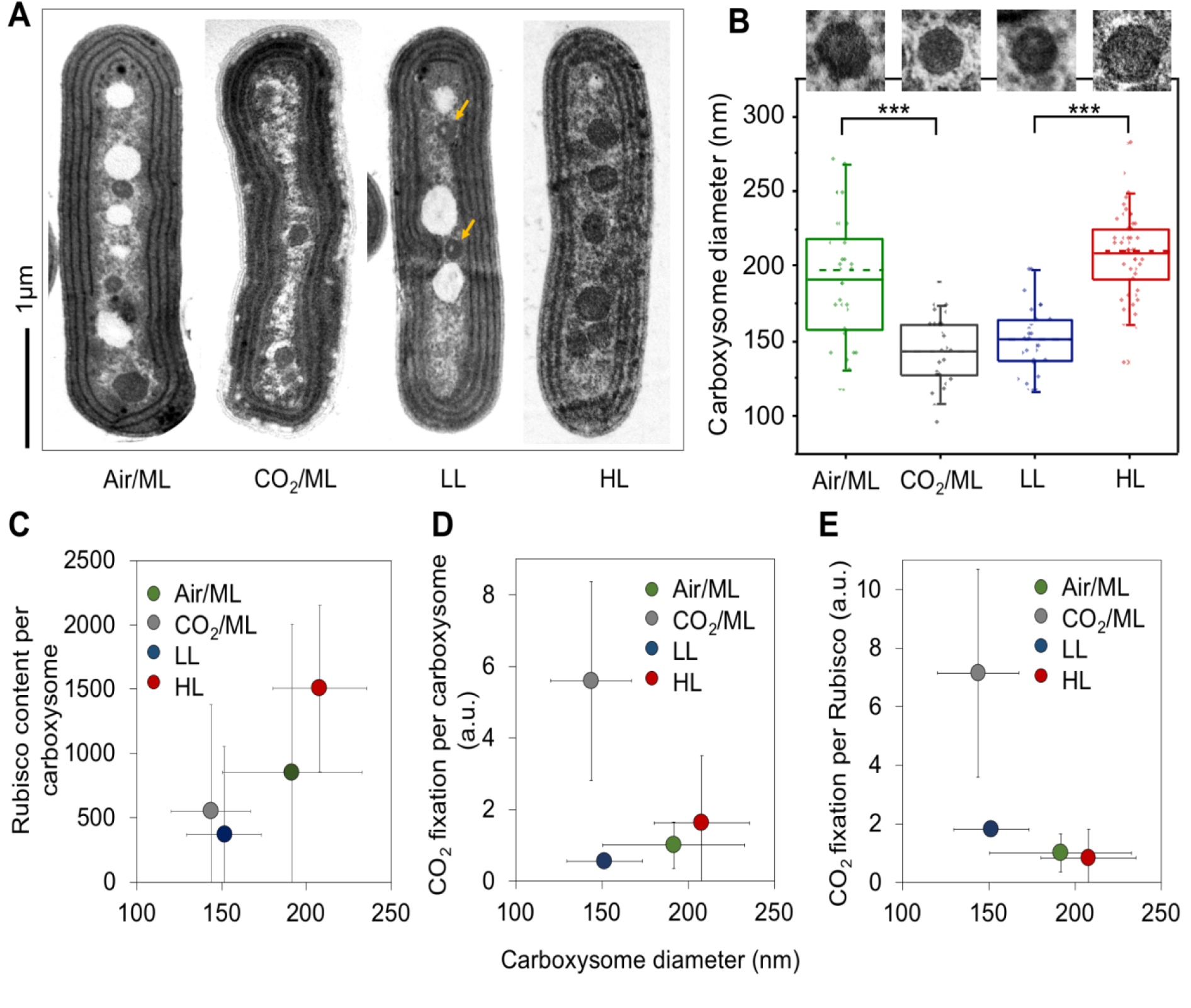
Variations of the carboxysome size and carbon fixation under Air/ML, CO_2_, LL and HL. **(A)** Thin-section electron microscopy (EM) images showing individual carboxysomes in the Syn7942 WT cells under Air/ML, CO_2_, LL and HL treatments Yellow arrows indicate the carboxysomes with spaces of low protein density under LL. More EM images are shown in Supplemental Figure 9. Scale bar indicates 1 μm. **(B)** Changes in the carboxysome diameter under Air/ML, CO_2_, LL and HL measured from EM (*n* = 33, 25, 27 and 51, respectively), with representative carboxysome images depicted above. Dashed lines indicate medians and solid lines indicate means. Differences in the carboxysome diameter are significant between CO_2_ and air (*p* = 1.92 × 10^−14^) and between LL and HL (*p* = 8.29 × 10^−7^), indicated as ***. **(C)** Correlation between the carboxysome size and the Rubisco content per carboxysome under Air/ML, CO_2_, LL and HL. **(D)** Correlation between the carboxysome size and CO_2_ fixation per carboxysome. **(E)** Correlation between the carboxysome size and CO_2_ fixation per Rubisco of the carboxysomes. Carboxysome diameters and CO_2_ fixation are presented as average ± SD, whereas the carboxysome total protein content and Rubisco content are shown as Peak value ± HWHM.

We also find that CO_2_-fixing activity per carboxysome increases as the β-carboxysome structure enlarges, which is correlated to strong light intensity during cell growth (Figure 4D), demonstrating the correlation between β-carboxysome structure and function *in vivo*. Moreover, under HL the CO_2_-fixation activity per Rubisco of the β-carboxysome declines as the carboxysome size and Rubisco density in the carboxysome lumen increase (Figure 4E, Supplemental Table 5). This may suggest that Rubisco density and local Rubisco packing are important for determining CO_2_-fixation activity of individual Rubisco (Supplemental Table 5). Interestingly, the relatively small β-carboxysomes under 3% CO_2_ exhibit high CO_2_-fixing activities per Rubisco and per carboxysome, compared with β-carboxysomes under other conditions. The enhanced carbon fixation capacity under 3% CO_2_ might be correlated with the increase in CcmK3 and CcmK4 content (Figure 3A, Table 1), as it has been shown that depletion of CcmK3/CcmK4 impedes carbon fixation of carboxysomes (Rae et al., 2012).

### Patterns of spatial localization and diffusion of β-carboxysomes in live cells change dynamically depending upon light intensity during growth

The patterns of β-carboxysome localization within the cyanobacterial cells appears to be crucial for carboxysome biogenesis and metabolic function (Savage et al., 2010; Sun et al., 2016). We measured the organizational dynamics of β-carboxysomes with distinct diameters in Syn7942 under different light intensities, using time-lapse confocal fluorescence imaging on the RbcL-YFP Syn7942 strain. Previous studies have shown that tagging of RbcL with fluorescent proteins does not obstruct β-carboxysome assembly and function in Syn7942 (Savage et al., 2010; Cameron et al., 2013; Chen et al., 2013; Sun et al., 2016). During time-lapse confocal imaging, we applied illumination on the cell samples, similar to that used for cell growth, in order to maintain cell physiology. We find that the overall mobility of individual β-carboxysomes within cyanobacterial cells is non-Brownian (Figure 5A, Supplemental Movie 1). Carboxysomes under HL display larger diffusive regions than those under LL. The mean square displacement (MSD) of tracked carboxysomes increased with the rise of light intensity (Figure 5B), as did the mean microscopic diffusion coefficient of individual carboxysomes (Figure 5C): an average diffusion coefficient of 2.76 ± 2.83 × 10^−5^ µm^2^·s^−1^ for HL (mean ± SD, *n* = 105), 1.48 ± 1.03 × 10^−5^ µm^2^·s^−1^ for ML (*n* = 84), and 0.28 ± 0.19 × 10^−5^ µm^2^·s^−1^ for LL (*n* = 336). It is interesting that the mobility of carboxysomes does not exhibit typical constrained diffusion – asymptotic MSD values at higher values of 𝛕 (Robson et al., 2013) – but rather exhibits anomalous diffusion at higher values of 𝛕 characterized by a non-linear relation, which can be observed in the intracellular protein mobility traces of other cellular systems (Lenn et al., 2008; Wollman et al., 2017). These results indicate the intracellular restrictions, for example the proposed interactions with the cytoskeletal system (Savage et al., 2010), McdA and McdB (MacCready et al., 2018) and ParA-mediated chromosome segregation (Jain et al., 2012), may mediate carboxysome positioning, but do not completely confine the mobility of carboxysomes. Notably, carboxysomes with a larger diameter (Figure 4) generated under HL present a higher diffusion coefficient compares with carboxysomes with relatively smaller size under ML and LL. However, there is no apparent correlation between the diffusion coefficient of carboxysomes and their size in the same light conditions (Supplemental Figure 10).

**Figure 5.**
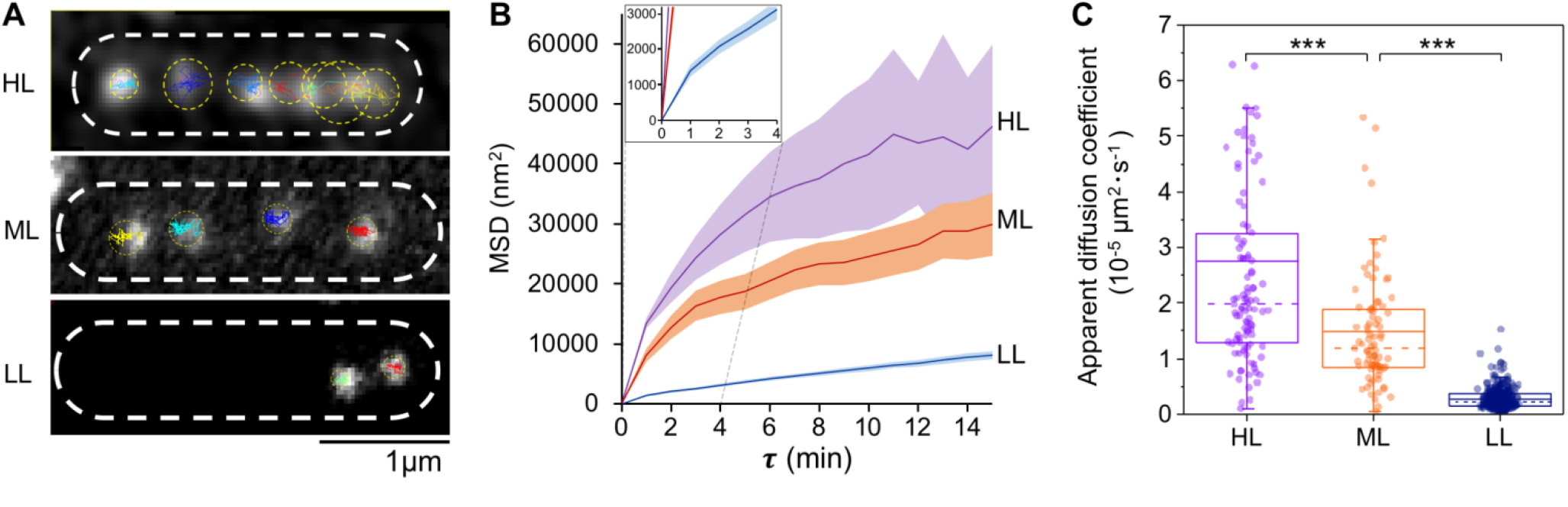
Spatial localization and diffusion dynamics of carboxysomes in Syn7942 cells are dependent on light intensity. **(A)** Tracking of carboxysome diffusion in cells grown under HL, ML and LL. Colored lines indicate the diffusion trajectories of each carboxysomes and circles represent the diffusion areas of each carboxysomes over 60 mins. Scale bar indicates 1 μm. **(B)** Non-linear MSD (Mean Square Displacement) *vs*. the time interval (𝛕) profiles suggest the mobility of carboxysomes in Syn7942 cells grown under HL, ML and LL. Inset, zoom-in view of the MSD profile under LL. **(C)** Diffusion coefficient of carboxysomes *in vivo* decreases significantly when the light intensity reduces: 2.76 ± 2.83 × 10^−5^ µm^2^·s^−1^ for HL (mean ± SD, *n* = 105), 1.48 ± 1.03 × 10^−5^ µm^2^·s^−1^ for ML (*n* = 84), and 0.28 ± 0.19 × 10^−5^ µm^2^·s^−1^ for LL (*n* = 336). *P* = 3.05 × 10^−5^ between HL and ML; *p* = 2.77 × 10^−5^ between ML and LL, two-tailed Student’s t-test).

## Discussion

Precise quantification of the protein stoichiometry and organizational regulation of carboxysomes provides insight into their assembly principles, structure and function. In this work, we functionally fused fluorescent protein tags to the building blocks in β-carboxysomes and exploited advanced “Physics of Life” technologies, in particular using bespoke single-molecule fluorescence microscopy to count the actual protein stoichiometry of β-carboxysomes in Syn7942 cells, at the single-organelle level. This approach minimizes the ensemble averaging encountered in bulk estimations from proteomic and immunoblot analysis. We characterized the stoichiometric flexibility of carboxysome proteins within individual polyhedral structures towards environmental variations. Variability of the protein stoichiometry and size of carboxysomes likely provides the structural foundation for the physiological regulation of carboxysome formation and carbon fixation activity. Given the shared structural features of carboxysomes and other BMCs, we believe that this work opens up new opportunities to quantitatively evaluate protein abundance and decipher the formation of all BMC organelles, in both native forms and synthetic variants.

Despite prior efforts on understanding carboxysome structure and function, the relative stoichiometry of functional carboxysome components in their native cell environment − key information required for reconstituting entire active carboxysome structures in synthetic biology (Fang et al., 2018), was still unclear. The major challenges have been the poor specificity of immunoblots and mass spectrometry, given the homology of carboxysome proteins and the lack of effective purification of intact carboxysomes from host cells, as well as the heterogeneity of carboxysome structures (Long et al., 2005). The previous model of carboxysome protein stoichiometry was based on the total amount of proteins in cell lysates (Long et al., 2011) and does not directly reflect the stoichiometry of carboxysome proteins in the organelle, given the possible free-standing carboxysome components in the cytosol (Dai et al., 2018). We have recently reported the isolation of β-carboxysomes from Syn7942 and the structural and mechanical exploration of the organelles (Faulkner et al., 2017). Interestingly, some components, i.e. CcmO, CcmN, CcmP and RbcX, were not detectable by mass spectrometry in the isolated carboxysomes, likely due to their low content or potential loss of carboxysome components during isolation. Here, as demonstrated, fluorescence tagging and Slimfield and confocal imaging enable single-organelle analysis of the protein stoichiometry of eight β-carboxysome proteins (including RbcX) and their regulation in their native context, and extends analyses of the assembly and action of carboxysomes. Microscopy imaging of fluorescently-tagged β-carboxysomes has been used to reveal their patterns of cellular localization, biogenesis pathways and light-dependent regulation in Syn7942 (Savage et al., 2010; Cameron et al., 2013; Chen et al., 2013; Sun et al., 2016; Niederhuber et al., 2017; MacCready et al., 2018). Although we cannot completely exclude the potential effects of YFP tags on carboxysome structure, we validate that YFP tagging to most of the structural components does not impede formation of functional carboxysome structures, suggesting the physiological relevance of the determined protein stoichiometry in the carboxysome in the presence of fluorescence tags. This flexibility emphasizes the extraordinary capacity of the carboxysome structure in adjusting their protein stoichiometry and accommodating foreign proteins while maintaining functionality, indicating the possibility of manipulating carboxysome organization in bioengineering for diverse purposes. Exceptionally, fluorescence tagging on CcmP and CcmO does not show normal carboxysome assembly and localization compared to other YFP-tagged strains (Supplemental Figure 11). In this work, therefore, we did not include estimation of the protein abundance of CcmP and CcmO, as well as RbcL and CcmK2 that cannot be fully tagged with YFP.

Numerous studies have described the regulation of carboxysome protein expression at the transcriptional level (McGinn et al., 2003; Woodger et al., 2003; Schwarz et al., 2011). Counting protein abundance of β-carboxysomes at different cell growth conditions enables direct characterization of the stoichiometric plasticity of carboxysome building components in the cells grown under not only the same environmental condition but also a range of various conditions (Figure 6A). Our observations elucidate the size variation of β-carboxysomes in Syn7942 cells grown under distinct environmental conditions (Figure 6B) and adjustable carbon fixation capacities of carboxysomes that may be closely linked to the protein organization and size of carboxysomes. Variations in the diameter of intact carboxysomes, ranging from 90 to 600 nm, have been also shown in previous studies not only in single species but also among distinct species (Shively et al., 1973; Price and Badger, 1991; Iancu et al., 2007; Liberton et al., 2011), suggesting the adaptation strategies exploited by cyanobacteria for regulating their CO_2_-fixing machines to survive in diverse niches. It may be related to the environment-sensitive protein-protein interactions that drive protein self-assembly and BMC formation (Faulkner et al., 2019). Moreover, the spatial positioning and mobility of β-carboxysomes in live cells appear to be independent of carboxysome diameter but show a strong dependence to light intensity, suggesting that light-dependent mechanisms might mediate carboxysome location and diffusion. Carboxysome spacing and partitioning have been suggested to be driven by different possible mechanisms, such as the cytoskeletal proteins ParA and MreB (Savage et al., 2010), ParA-mediated chromosome segregation (Jain et al., 2012) via filament-pull model (Ringgaard et al., 2009) or a diffusion-ratchet model (Vecchiarelli et al., 2013) as well as very recently the McdA and McdB that utilize a Brownian-ratchet mechanism to position carboxysomes (MacCready et al., 2018). Altogether, the organizational flexibility of β-carboxysomes, including modulatable protein stoichiometry, diameter and mobility, may represent the natural strategies for modifying shell permeability and enzyme encapsulation and ensuring structural and functional adaptations dependent on the local cellular environment.

**Figure 6.**
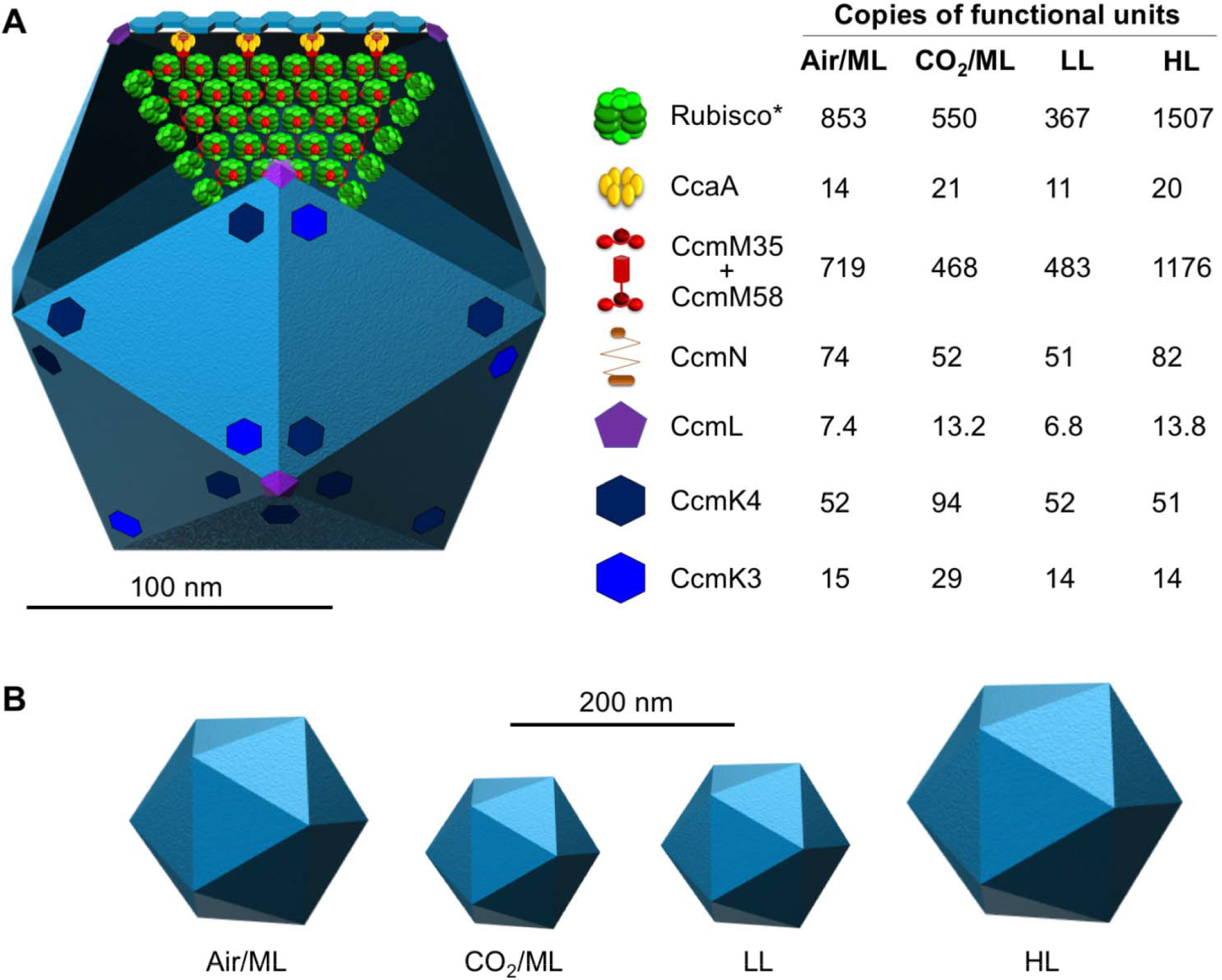
Model of the β-carboxysome structure and protein stoichiometry. **(A)** Diagram of an icosahedral carboxysome structure and organization of building components. The stoichiometry of each building component within the carboxysome and its variations in response to changes in CO_2_ and light intensity are shown on the right (See also Table 1). *Rubisco content was estimated from RbcS stoichiometry based on the RbcL_8_S_8_ Rubisco structure. The majority of shell facets shown in light blue is tiled by the major shell protein CcmK2. The total abundance of CcmM58 and CcmM35 was estimated. The components RbcL, CcmK2, CcmO and CcmP were not directly determined in this work and thus are not shown in this model. **(B)** The carboxysome diameter is variable in response to changes in the CO_2_ level and light intensity.

The estimated number of CcmL pentamers per carboxysome could be less than 12, demonstrating explicitly that it is not a prerequisite for CcmL pentamers to occupy all 12 vertices of the icosahedral shell to ensure complete formation of functional carboxysomes. This hypothesis has been validated by previous observations that BMC shells in the absence of pentamers have no significant morphological changes (Cai et al., 2009; Lassila et al., 2014; Hagen et al., 2018a). These “non-complete capped” forms appear to be prevalent among the resultant carboxysomes under Air/ML and LL (Figure 2C), unlike the procarboxysomes (Cameron et al., 2013) or “immature” carboxysomes which are incapable of establishing an oxidative microenvironment for cargo enzymes (Chen et al., 2013). Whether the loss of capping CcmL will create large space within the shell, as a possible mechanism of modulating shell permeability, or will be compensated for by incorporation of other shell proteins, for example the additional CcmP trimers that are speculated to be responsible for permeability, remains to be further investigated. Our results also suggest that carboxysomes could possess a flexible molecular architecture, resonating with the observation of structural “breathing” of virus capsids which has been reported to be key to cope with temperature change (Roivainen et al., 1993; Li et al., 1994). Carboxysomes, though structurally resembling virus capsids, have been shown to be mechanically softer than the P22 virus capsid by a factor of ∼10, suggesting greater flexibility of protein-protein interactions within the carboxysome structure (Faulkner et al., 2017). The capping flexibility of pentamers may represent the dynamic nature of shell assembly probably in the second timescale and tunable protein-protein interactions in the shell, as characterized recently (Sutter et al., 2016; Faulkner et al., 2019).

It was proposed that CcmM58 proteins are confined to a subshell layer for linking Rubisco, CcaA and CcmN to the shell, whereas CcmM35 molecules are predominantly located in the core to stimulate Rubisco aggregation (Rae et al., 2013). A recent study revealed that CcmM35 and CcmM58 display similar distribution profiles in carboxysomes and are both integrated within the core of the carboxysome (Niederhuber et al., 2017). Fluorescence tagging at the protein C-terminus exploited in this work allowed us to only estimate the total amounts of CcmM but not distinguish CcmM35 and CcmM58, which can be addressed by N-terminal labeling of CcmM58 in our future study. Compared with the previous model that was based on protein stoichiometry of cell lysates (Long et al., 2011), our relative quantifications determined under the Air/ML condition show the 4.9-fold and 2.2-fold increases in the ratios of Rubisco/CcmM and Rubisco/CcaA, respectively (Figure 6A, Supplemental Table 5). The discrepancy may be caused by different sampling methods and cultivation conditions.

Based on immunoblot analysis of cell lysates, the previous model has proposed an imbalanced ratio of RbcL to RbcS (∼8:5), likely due to the binding of CcmM to Rubisco replacing 3 RbcS subunits (Long et al., 2011). This result was similar to our immunoblot quantification from cell lysates (Supplemental Table 2). Recent studies indicate that CcmM interacts with Rubisco (RbcL_8_S_8_) at distinct sites, without displacing RbcS (Ryan et al., 2018; Wang et al., 2019). Based on the L_8_S_8_ ratio and RbcS abundance per carboxysome determined, we estimate that there are approximately 853, 550, 367, and 1507 copies of Rubisco per β-carboxysome under Air/ML, CO_2_/ML, LL, and HL, respectively (Figure 6A, Table 1). Even the lowest Rubisco abundance per β-carboxysome (an average diameter of 151 nm) under LL is still greater than the Rubisco abundance per α-carboxysome (an average diameter of 123 nm) (Iancu et al., 2007) by a factor of 1.6. This finding confirms the different interior organization of the two classes of carboxysomes: densely packed with Rubisco forming paracrystalline arrays inside the β-carboxysome (Faulkner et al., 2017) and random packing of Rubisco in the α-carboxysome (Iancu et al., 2007; Iancu et al., 2010). The different interior structures may be ascribed to their distinct biogenesis pathways: biogenesis of β-carboxysomes is initiated from the nucleation of Rubisco and CcmM35 and then the shell encapsulation (Cameron et al., 2013); whereas α-carboxysome assembly appears to start from shell formation (Menon et al., 2008) or a simultaneous shell-interior assembly (Iancu et al., 2010).

While the abundance of most of the structural components varies, the ratio of CcmK4 and CcmK3 is relatively unaffected (ranging from 3.6 to 4.1, Supplemental Table 5) under the tested growth conditions, implying their spatial colocalization within the carboxysome shell (Figure 6A). This is reminiscent of the recent observation that CcmK3 and CcmK4 can form a heterohexameric complex with a 1:2 stoichiometry and further form dodecamers in a pH-dependent manner (Sommer et al., 2019). The *ccmK3* and *ccmK4* genes are located in the same operon that is distant from the *ccm* operon and they may have different expression regulation compared with other carboxysome components (Rae et al., 2012; Sommer et al., 2017). The balanced expression and structural cooperation of CcmK3 and CcmK4 may be crucial for the fine-tuning of carboxysome permeability towards environmental stress.

Rational design, construction and modulation of bioinspired materials with structural and functional integrity are the major challenges in synthetic biology and protein engineering. Given their self-assembly, modularity and high efficiency in enhancing carbon fixation, carboxysomes have attracted tremendous interest to engineering this CO_2_-fixing organelle into other organisms, for example C_3_ plants, with the intent of increasing photosynthetic efficiency and crop production (Lin et al., 2014b; Lin et al., 2014a; Occhialini et al., 2016; Long et al., 2018). Recently, we have reported the engineering of functional β-carboxysome structures in *E. coli* – a step towards constructing functional β-carboxysomes in eukaryotic organisms (Fang et al., 2018). Our present study, by evaluating the actual protein stoichiometry and structural variability of native β-carboxysomes, sheds light on the molecular basis underlying the assembly, formation and regulation of functional carboxysomes. It will empower bioengineering to construct BMC-based nano-bioreactors and scaffolds, with functional and tunable compositions and architectures, for metabolic reprogramming and targeted synthetic molecular delivery. A deeper understanding of carboxysome structure and the developed imaging techniques will be broadly extended to other BMCs and macromolecular systems.

## Materials and Methods

### Bacterial strains, growth conditions, light and CO_2_ treatment, and generation of mutants

Wild-type (WT) and mutant *Synechococcus elongatus* PCC7942 (Syn7942) strains were grown in BG-11 medium in culture flasks with constant shaking or on BG-11 plates containing 1.5% (w/v) agar at 30°C. Syn7942 WT and mutants were maintained and grown under different intensities of constant white LED light illumination: 80 μE·m^−2^·s^−1^ as HL (higher light in ambient air), 50 μE·m^−2^·s^−1^ as Air/ML (moderate light in ambient air), 10 μE·m^−2^·s^−1^ as LL (lower light in ambient air) to ensure full acclimation, respectively. Cultures were grown in air without an additional CO_2_ source, except for the CO_2_ treatment experiment in which Syn7942 cultures in the growth incubators were aerated with 3% CO_2_ under moderate light (CO_2_/ML).

Cultures were constantly diluted with fresh medium to maintain exponential growth phase for the following imaging and biochemical analysis. *Escherichia coli* strains used in this work, DH5a and BW25113, were grown aerobically at 30 or 37°C in Luria-Broth medium. Medium supplements were used, where appropriate, at the following final concentrations: ampicillin 100 mg·mL^−1^, chloramphenicol 10 mg·mL^−1^, apramycin 50 mg·mL^−1^, and arabinose 100 mM.

All YFP-fusion mutants were generated following the REDIRECT protocol (Supplemental Figure 1) (Gust et al., 2002), by inserting the *eyfp*:*apramycin* DNA fragment to the C-terminus of individual carboxysome genes based on homologous recombination (Supplemental Table 6). Primers used in this work were listed in Supplemental Table 7. The same strategy was also applied for the mYPet mutant. For these mutant strains, BG-11 medium was supplemented with apramycin at 50 μg·mL^−1^.

### Cell doubling time and growth curve measurement

Cultures were inoculated at OD_750_ of 0.05-0.1 with fresh BG-11. Growth of cells was monitored at OD_750_ using a spectrophotometer (Jenway 6300 spectrophotometer, Jenway, UK) every 24 hours. Doubling times were calculated using exponential phase of growth from day 1 to day 4. Four biological replicates from different culture flasks were recorded. Data are presented as mean ± standard deviation (SD). For each experiment, at least three biological replicates from different culture flasks were analyzed.

### Slimfield microscopy and data analysis

Live cells were applied at the small volume onto the BG-11 agarose pad at 0.25 mm thickness to maintain physiological growth, air dried to remove excessive medium and then assembled with plasma cleaned (Harrick-Plasma) glass cover slips. A dual-color bespoke laser excitation single-molecule fluorescence microscope was used utilizing narrow epifluorescence excitation of 10 µm full width at half maximum (FWHM) in the sample plane to generate Slimfield illumination using narrowfield epifluorescence (Wollman and Leake, 2016; Wollman et al., 2016b; Wollman et al., 2017). This was incident on a sample mounted on a Mad City Labs nanostage built on an inverted Zeiss microscope body consisting of a 20 mW 514 nm wavelength laser. A Chroma GFP/mCherry dichroic was mounted under the Olympus 100x NA = 1.49 TIRF (total internal reflection fluorescence) objective, which delivers 10 mW excitation power. The image was split into YFP and chlorophyll channels using a bespoke color splitter utilizing a Chroma dichroic split at 560 nm with 542 nm and 600 nm, 25 nm bandwidth filters. Imaging was done with an Andor iXon 128 × 128 pixel EMCCD camera (iXon DV860-BI, Andor Technology, UK), at a pixel magnification of 80 nm/pixel using 5 ms camera exposure time. Excitation intensity was initially reduced by 100x using and ND = 2 or 1 attenuation filter for high copy number strains (all except CcmL and RbcX) to avoid pixel saturation on the EMCCD camera detector before a full-power photobleaching. Sample sizes for individual strains are 60 (RbcS), 219 (CcmK3), 77 (CcmK4), 316 (CcmL), 71 (CcmM), 86 (CcmN), 95 (CcaA) and 211 (RbcX), respectively. Each population of carboxysomes comes from 20-30 fields of view, with 1-7 cells per field of view.

The analysis was performed using bespoke MATLAB (Mathworks) software (Miller et al., 2015) with previously outlined methods (Llorente-Garcia et al., 2014; Wollman et al., 2016a; Beattie et al., 2017; Lund et al., 2018; Stracy et al., 2018). In brief, candidate bright fluorescent foci were identified in images using morphological transformation and thresholding. The sub-pixel centroids of these foci were determined using iterative Gaussian masking and their intensity quantified as the summed intensity inside a 5-pixel radius region of interest (ROI) corrected for the mean background intensity inside a surrounding 17 × 17 pixel ROI (Delalez et al., 2010; Leake, 2014). Foci were accepted and tracked through time if they had a signal-to-noise ratio, defined as the mean intensity in the circular ROI divided by the standard deviation in the outer ROI, over 0.4. The characteristic intensity of single YFP/mYPet was measured from the distribution of detected foci intensity towards the end of the photobleaching (Figure 1), confirmed by comparing the obtained value to individual photobleaching steps obtained using edge-preserving filtration (Figure 1) (Leake et al., 2003; Leake et al., 2004). The stoichiometry of foci was then determined through cell-by-cell based Slimfield imaging using numerical integration of pixel intensities (Wollman and Leake, 2015) in each carboxysome divided by the intensity of a single YFP (Figure 1B).

For high-copy-number strains, intensity of carboxysomes was very high compared to the chlorophyll but for CcmL (typically ∼2x, compare Supplemental Figure 3 with Supplemental Figure 4A) the fluorescence intensity per carboxysome was comparable (although generally brighter) to small regions of bright chlorophyll, detected as foci by our software, as confirmed by looking at the parental strain with no YFP present. To correct for this chlorophyll content, we tracked parental WT Syn7942 cells as YFP-labelled cells to calculate the apparent chlorophyll stoichiometry distribution (Supplemental Figure 4A). The CcmL distribution was then corrected by subtracting the apparent chlorophyll distribution. To investigate putative periodic features in the stoichiometry distribution, we used the raw uncorrected values to minimize dephasing artefacts (Figure 4C) using a kernel width of 0.5 molecules (equivalent to the error in determining the characteristic intensity). The peak values in other strains were far from the chlorophyll peak and so unaffected by this correction.

### Confocal microscopy imaging and data analysis

Preparation of Syn7942 cells for confocal microscopy was performed as described earlier (Liu et al., 2012; Casella et al., 2017). Cells were maintained under different growth conditions prior to microscopy imaging, to ensure full acclimation. Confocal fluorescence images (12-bit, 512 × 512 pixels) were recorded using a Zeiss LSM780 with an alpha Plan-Fluor 100x oil immersion objective (NA 1.45) and excitation at 514 nm from an Argon laser. YFP and chlorophyll fluorescence were captured at 520−550 nm and 660−700 nm, respectively. The image pixel size was 41.5 nm. The pixel dwell time was 0.64 μs and the frame averaging was 8, resulting in an effective frame time of ∼1.5 s. The pinhole was set to give *z* axis resolution of 1 μm. Live-cell confocal fluorescence images were recorded from at least five different cultures. The sample stage was pre-incubated and thermo-controlled at 30°C before and during imaging. Zoom settings were set to have each carboxysome visualized with a minimum of 8 × 8 pixels array to allow sufficient profiling of carboxysome signals by peak intensity recognition and measurement. All images were captured with all pixels below saturation.

Confocal microscopic images were processed using FIJI Trackmate plugins (Tinevez et al., 2017) to retrieve peak intensities of carboxysomes based on the Find Maxima detection algorithm. Noise tolerance was determined by background intensities in empty regions. Imaging for different treatments in the same strain was performed under the same imaging settings. For strains with visible cytosolic signals, the cytosolic background intensity was determined by the average peak intensities in non-carboxysome regions over the central line of the cell and was subtracted to obtain peak intensities. Raw data were processed by Origin Lab and MATLAB (Mathworks) for profile extraction and statistical analysis and the goodness-of-fit parameter for Violin plot visualization. Violin plots were generated by R to illustrate the fluorescence intensity distribution of individual building proteins per carboxysome fitted by kernel smooth fitting. The representative values and deviations of signal intensities were represented by Peak value ± half width at half maximum (HWHM) measured from kernel density fitted profiles, respectively. The significance of differences between treatments was evaluated by Mann-Whitney U-tests pair-wisely (Supplemental Table 4). Standard errors of sampling were determined through randomized grouping of intensity entries, with each group containing a minimum of 70–100 entries. Errors were controlled below 5% to have accurate estimation from the distributions. The relative protein abundance of carboxysomes was estimated by confocal imaging under Air/ML, CO_2_/ML, LL, and HL was normalized by the definite copy number of each strain under Air/ML determined by Slimfield imaging.

### Live-cell time-lapse confocal imaging and data analysis

A 2 mm-thick BG-11 agar mat was prepared in stacked sandwiches to accommodate drops of diluted Syn7942 cells. Cells were incubated on the BG-11 agar mat on the microscope for 1-2 hours before imaging. The continuous light illumination was provided at the intensity relatively equal to HL, ML, or LL that were used for cell growth, in order to maintain cell physiology. The same illumination was applied to the cells during time-lapse imaging with a hand-made module that switched off the light during laser scanning (less than 5 s per minute intervals). The interval time was set to 60 s to guarantee sufficient light illumination between imaging. The laser power was set to the minimum (1%) to reduce the bleaching for signals during long-term tracking. Images were initially corrected for horizontal drifting by Descriptor-based series registration (2d/3d+T) plugin, and then were processed by the Trackmate plugin in FIJI for particle tracking. Retrieved track data was analyzed using bespoke MATLAB (Mathworks) scripts for MSD. Diffusion coefficient calculations and data visualization were modified as previously described (Ewers et al., 2005; Sbalzarini and Koumoutsakos, 2005). Diffusion coefficients were calculated by fitting the first 6 points of the MSD *vs*. 𝝉 curves. As the MSD *vs*. 𝝉 curves indicated potentially non-Brownian diffusion at higher 𝝉 values, we described the diffusion coefficients as “apparent diffusion coefficients”. Tracking and diffusion coefficient determination were tested by computational simulations (Supplemental Movie 2). Bespoke Matlab code was written to generate simulated image stacks of carboxysomes diffusing inside cells. Images were simulated by integrating a model 3D point spread function over a 3D model for the cell structure (Wollman and Leake 2015). This model comprises an inner cytosol surrounded by thylakoid membranes (indicated by chlorophyll fluorescence) and 3 carboxysomes with a diameter of 200 nm. Each component’s intensity was adjusted to match real images before representative Poisson noise was applied. Carboxysomes were simulated undergoing Brownian motion with a diffusion coefficient of 1.3 × 10^−5^ µm^2^·s^−1^ over 40 image frames. Trackmate tracking and diffusion coefficient calculation yielded a mean diffusion coefficient of 1.32 ± 0.02 × 10^−5^ µm^2^·s^−1^, giving a 1.5% error.

### Immunoblot analysis

Immunoblot examination was carried out following the procedure described previously (Sun et al., 2016). 150 µg of cell lysate, measured by Pierce Coomassie (Bradford) Protein Assay Kit (Thermo Fisher Scientific), was loaded on 10% (v/v) denaturing SDS-PAGE gels. Immunoblot analysis was performed using the primary mouse monoclonal anti-GFP (Invitrogen, 33-2600), capable of recognizing series of GFP variants including YFP, the rabbit polyclonal anti-RbcL (Agrisera, AS03 037), the horseradish peroxidase-conjugated goat anti-mouse IgG secondary antibody (Promega, W4021) and a Goat anti-Rabbit IgG (H&L), HRP conjugated (Agrisera AS10 1461). Anti-CcmK2 antibody was kindly provided by the Kerfeld lab (Michigan State University, US) (Cai et al., 2016). Protein quantification from immunoblot data was carried out using FIJI. Our nominal assumption that the ratios of YFP-tagged to total RbcL or CcmK2 in carboxysomes are similar to those in cell lysates.

### *In vivo* carbon fixation assay

*In vivo* carbon fixation assay was carried out to determine carbon fixation of Syn7942 WT and mutant cells, as described in the previous work (Sun et al., 2016). For each WT and mutant, at least three biological replicates from different culture flasks were assayed. Significance was assessed by two-tailed Student’s t-tests.

### Electron microscopy and carboxysome size measurement

Electron microscopy was carried out as described previously (Liu et al., 2008; Sun et al., 2016). Carboxysome diameter was measured as described previously (Faulkner et al., 2017) and was analyzed using Origin.

## Accession Numbers

Accession numbers of genes in this article are provided in Supplemental Table 6.

## Supplemental Data

**Supplemental Figure 1.** Construction and verification of Syn7942 strains with YFP fusion to individual carboxysome proteins.

**Supplemental Figure 2.** Immunoblot analysis of the YFP-tagged Syn7942 strains using the anti-GFP, anti-RbcL and anti-CcmK antibodies of soluble fractions in this study based on SDS-PAGE.

**Supplemental Figure 3.** Slimfield images of YFP-fusion cells under Air/ML and stoichiometric histogram of copies of YFP molecules per carboxysome.

**Supplemental Figure 4.** Normalization of chlorophyll during Slimfield imaging for Syn7942 strains.

**Supplemental Figure 5.** Comparison of YFP and mYPet tagging to RbcL reveals no differences in carboxysome localization, cell growth and carbon fixation, suggesting that there are no measurable artefacts due to putative effects of dimerization of the YFP tag.

**Supplemental Figure 6.** Confocal images of YFP-tagged cells.

**Supplemental Figure 7.** Confocal images of RbcS-YFP, CcmM-YFP, CcmK4-YFP and CcmK3-YFP cells under Air/ML, CO_2_, LL, and HL and distribution profiles of carboxysome protein signal intensity.

**Supplemental Figure 8.** Confocal images of CcmL-YFP, CcmN-YFP, CcaA-YFP and RbcX-YFP cells under Air/ML, CO_2_, LL, and HL and distribution profiles of carboxysome protein signal intensity (continuing Supplemental Figure 7).

**Supplemental Figure 9.** Thin-section EM images of WT Syn7942 cells under Air/ML, CO_2_/ML, LL and HL.

**Supplemental Figure 10.** Changes in the diffusion coefficient of carboxysomes in Syn7942 cells under HL, ML and LL are not dependent on the carboxysome size.

**Supplemental Figure 11.** CcmP-YFP and CcmO-YFP Syn7942 cells.

**Supplemental Table 1.** Cell growth, carbon fixation and cell dimensions of Syn7942 WT and YFP-fusion mutants under Air/ML.

**Supplemental Table 2.** Immunoblotting estimation of the stoichiometry of carboxysomal proteins in cell lysates.

**Supplemental Table 3.** Carboxysome content per cell under Air/ML, CO_2_/ML, LL and HL determined by confocal imaging.

**Supplemental Table 4.** Evaluation and quality control of quantitative microscopy.

**Supplemental Table 5.** Carboxysome properties in Syn7942 vary under Air/ML, CO_2_/ML, LL and HL, determined by Slimfield, confocal and EM imaging.

**Supplemental Table 6.** Accession numbers for genes/proteins in this work.

**Supplemental Table 7.** PCR primers used in this study for gene cloning and sequencing.

**Supplemental Movie 1.** Time-lapse confocal imaging reveals different diffusion dynamics of carboxysomes in the RbcL-YFP Syn7942 cells grown under HL, ML and LL conditions.

**Supplemental Movie 2.** Simulations of diffusing carboxysomes *in cellulo* validate tracking and diffusion coefficient determination.

## Acknowledgements

We thank Gregory F Dykes, Selene Casella and Alison Beckett for technical support of electron microscopy. We thank David Mason in confocal image analysis. We thank the Liverpool Centre for Cell Imaging for technical assistance and provision. Y.S., F.H. and L.-N.L. were supported by Royal Society (UF120411, IE131399, RGF\EA\180233, RGF\EA\181061 and URF\R\180030, L.-N.L.) and Biotechnology and Biological Sciences Research Council Grant (BB/M024202/1 and BB/R003890/1, L.-N.L.), Leverhulme Trust (ECF-2016-778, F.H.) and China Scholarship Council (Y.S.). M.L. was supported by a Medical Research Council grant (MR/K01580X/1), Biotechnology and Biological Sciences Research Council Grant (BB/N006453/1). A.J.M.W. was part-funded by the Wellcome Trust (204829) through Centre for Future Health at the University of York.

## Author contributions

L.-N.L and M.C.L designed research; Y.Q. A.J.M.W and F.H. performed research and analyzed data; L.-N.L, Y.Q., M.C.L., and A.J.M.W. wrote the paper.

## Competing interests

The authors declare no conflict of interest.

## Notes

#### Summary of Updates

This manuscript has been published in The Plant Cell: http://www.plantcell.org/content/early/2019/05/02/tpc.18.00787

