## Supplemental Information for "Single-Organelle Quantification Reveals Stoichiometric and Structural Variability of Carboxysomes Dependent on the Environment" (from another version of this preprint: version 1, 2019-03-05)

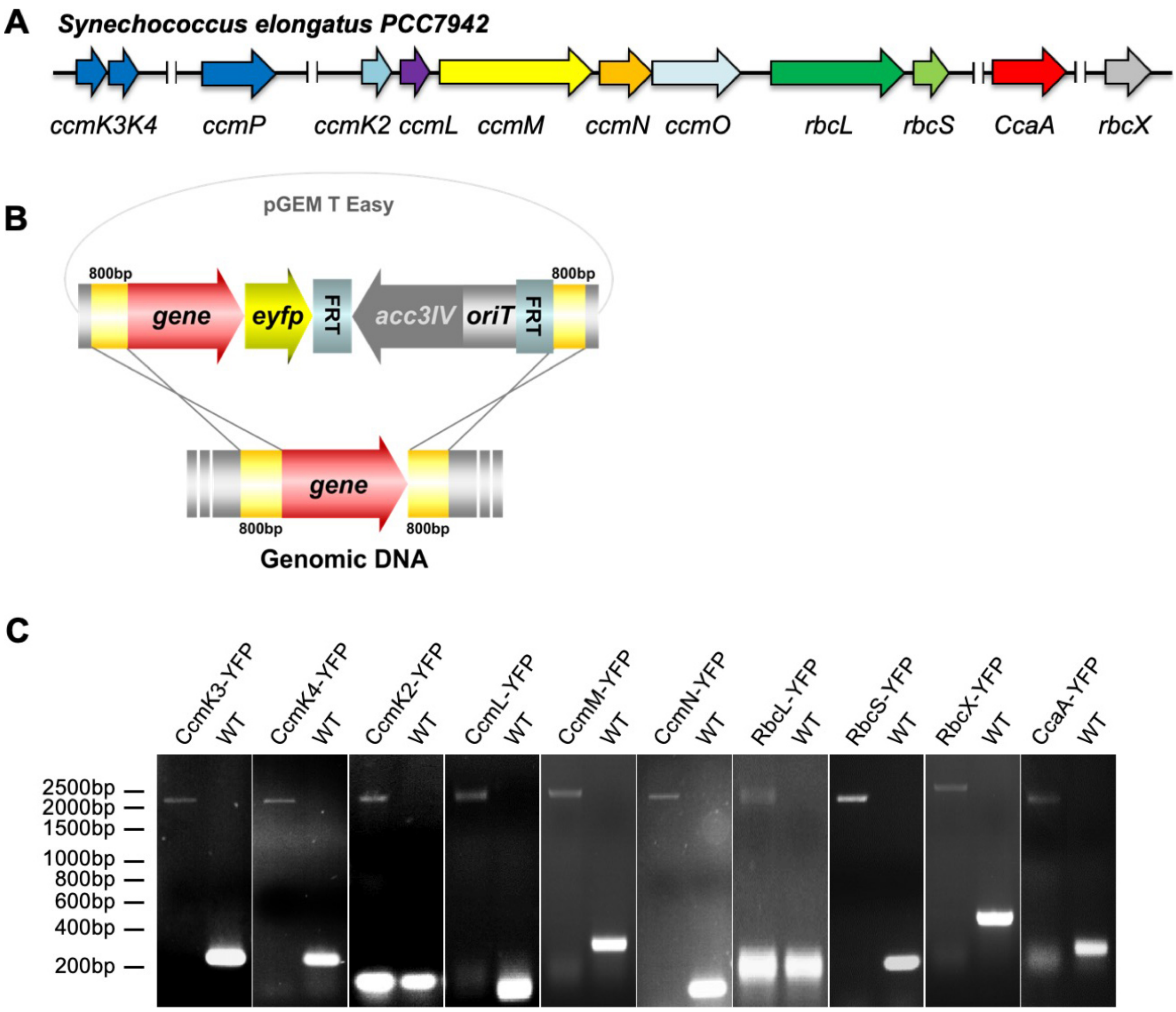


**Supplemental Figure 1. Construction and verification of Syn7942 strains with YFP fusion to individual carboxysome proteins.**
**(A)** Locations of the genes encoding carboxysome proteins in WT Syn7942.
**(B)** The strategy of YFP fusion using REDIRECT protocol.
**(C)** PCR verification of the full segregation of YFP-fusion strains, except for CcmK2-YFP and RbcL-YFP which are partially segregated (See also Supplemental Figure 2).

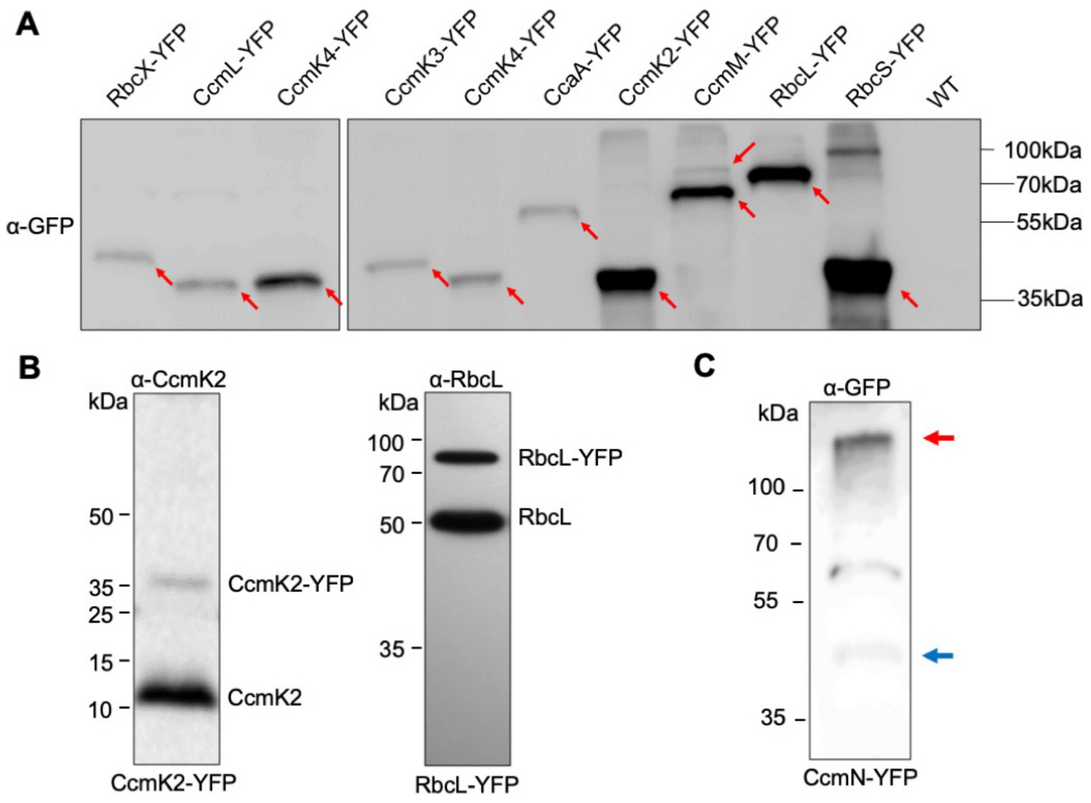


**Supplemental Figure 2. Immunoblot analysis of the YFP-tagged Syn7942 strains using the anti-GFP, anti-RbcL and anti-CcmK antibodies of soluble fractions in this study based on SDS-PAGE.**

**(A)** Immunoblot of soluble fractions of YFP-tagged strains using an anti-fluorescent protein antibody shows tagging of YFP to each carboxysome protein (red arrows). Note that CcmM-YFP shows two bands at 85 and 62 kDa, corresponding to CcmM58 and CcmM35, respectively. Samples with the same total protein concentration were loaded to relatively quantify the carboxysome protein content in the soluble fraction (See also Table 1).

**(B)** Immunoblot of soluble fractions of the partially-segregated CcmK2-YFP and RbcL-YFP strains, using anti-CcmK2 and anti-RbcL antibodies. It indicates that 29.2 ± 7.1 % ($n$ = 4) of total RbcL and 6.0 ± 0.7 % ($n$ = 3) of total CcmK2 proteins were tagged with YFP.

**(C)** Immunoblot of soluble fractions of the CcmN-YFP strain using an anti-fluorescent protein antibody, showing the presence of large aggregates of CcmN that failed to run into the gel, marked by the red arrow. Minor fractions of CcmN that run into the gel are marked by the blue arrow. For all western results, at least 3 independent experiments were done for statistical analysis.

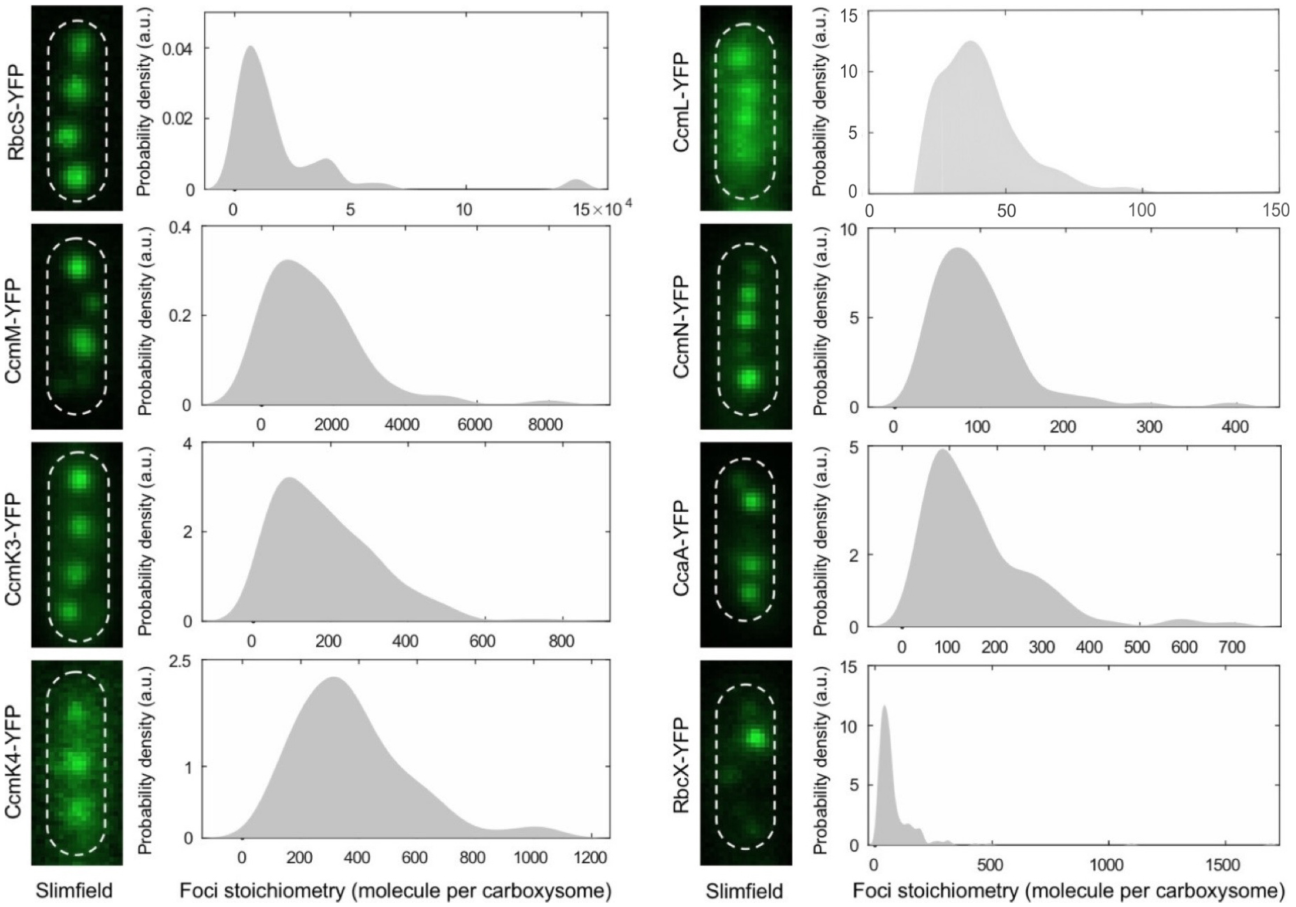


**Supplemental Figure 3. Slimfield images of YFP-fusion cells under Air/ML and stoichiometric histogram of copies of YFP molecules per carboxysome.**
Fluorescence foci (green) indicate individual carboxysomes in cells. Cell borders are outlined by white dash lines. In the histograms, the major peaks were acquired by kernel density fitting, representing the copy numbers of each proteins per carboxysome. Sample sizes for individual strains are 60 (RbcS), 219 (CcmK3), 77 (CcmK4), 316 (CcmL), 71 (CcmM), 86 (CcmN), 95 (CcaA) and 211 (RbcX), respectively.

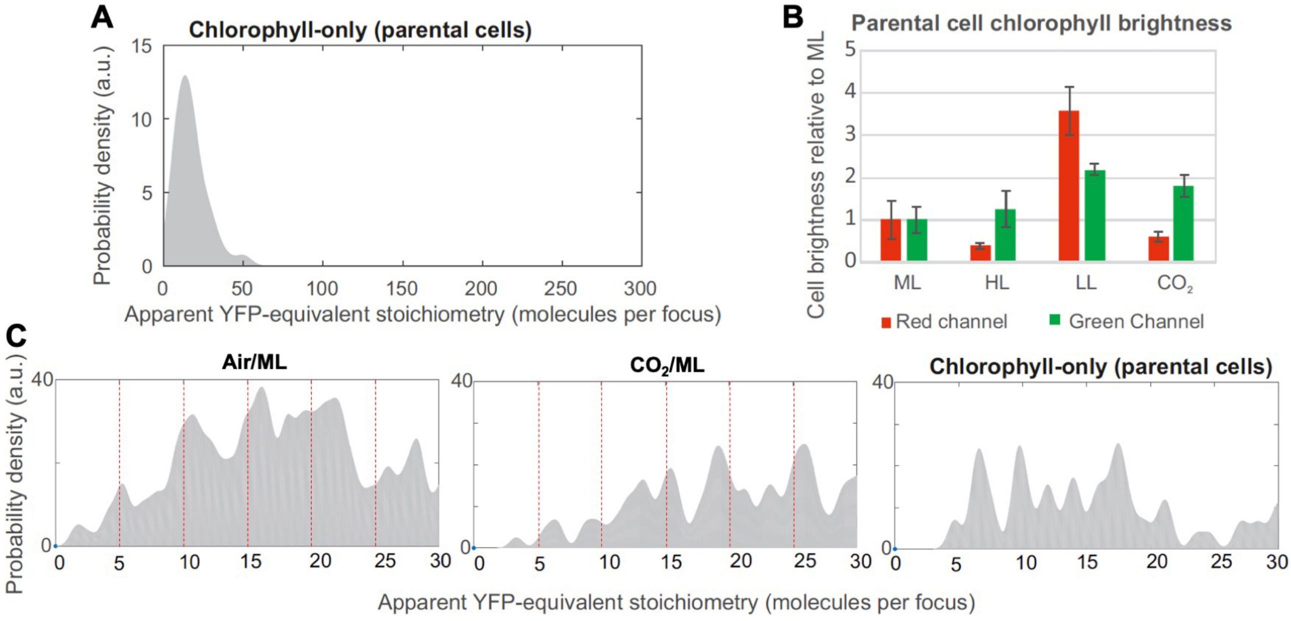


**Supplemental Figure 4. Normalization of chlorophyll during Slimfield imaging for Syn7942 strains.**

**(A)** The apparent YFP-equivalent stoichiometry of Chlorophyll estimated from Wild-type Syn7942 at Air/ML condition.

**(B)** Relative brightness of chlorophyll detected by Slimfield imaging in the YFP (green) channel and chlorophyll (red) channel, showing expected trends between growth conditions in the red channel but no clear trend in the green channel, such that the Air/ML distribution in **(A)** was used to correct all growth conditions shown in Figure 2.

**(C)** Uncorrected kernel density estimates for CcmL stoichiometries in Air/ML and $CO_2$/ML growth conditions and apparent stoichiometry of chlorophyll, generated with a kernel width of 0.5 YFPs. This low kernel width allows periodic features to be seen, in this case showing 5-mer periodicity in CcmL stoichiometries.

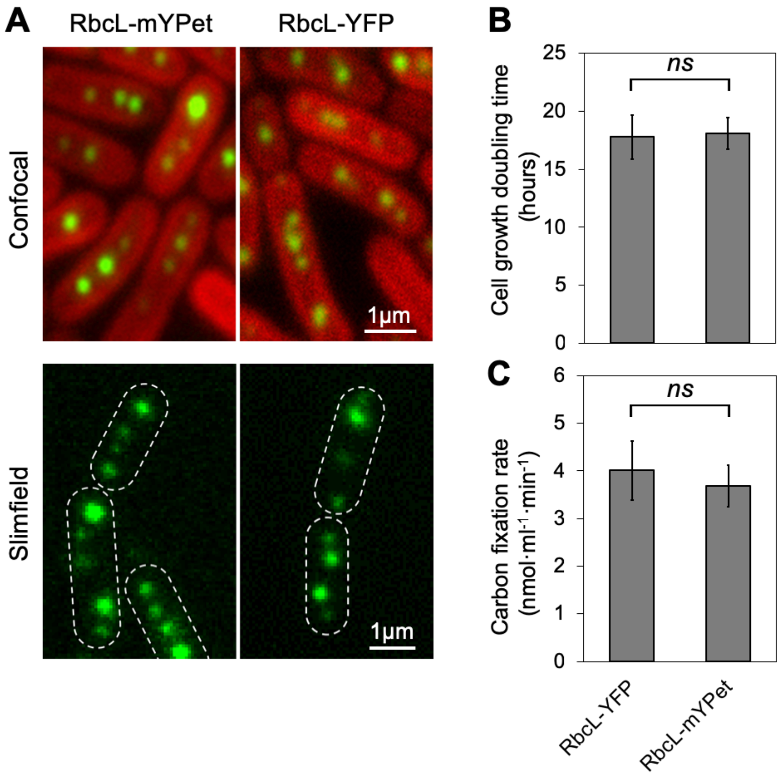


**Supplemental Figure 5. Comparison of YFP and mYPet tagging to RbcL reveals no differences in carboxysome localization, cell growth and carbon fixation, suggesting that there are no measurable artefacts due to putative effects of dimerization of the YFP tag.**
**(A)** Confocal and Slimfield microscopy images for RbcL-mYPet and RbcL-YFP. Scale bars indicate 1 μm.
**(B-C)** No significant effects were detected on the doubling time ($p$ = 0.82, $n$ = 4, Student's t-test) and carbon fixation capacity ($p$ = 0.50, $n$ = 3) of RbcL-mYPet and RbcL-YFP strains.

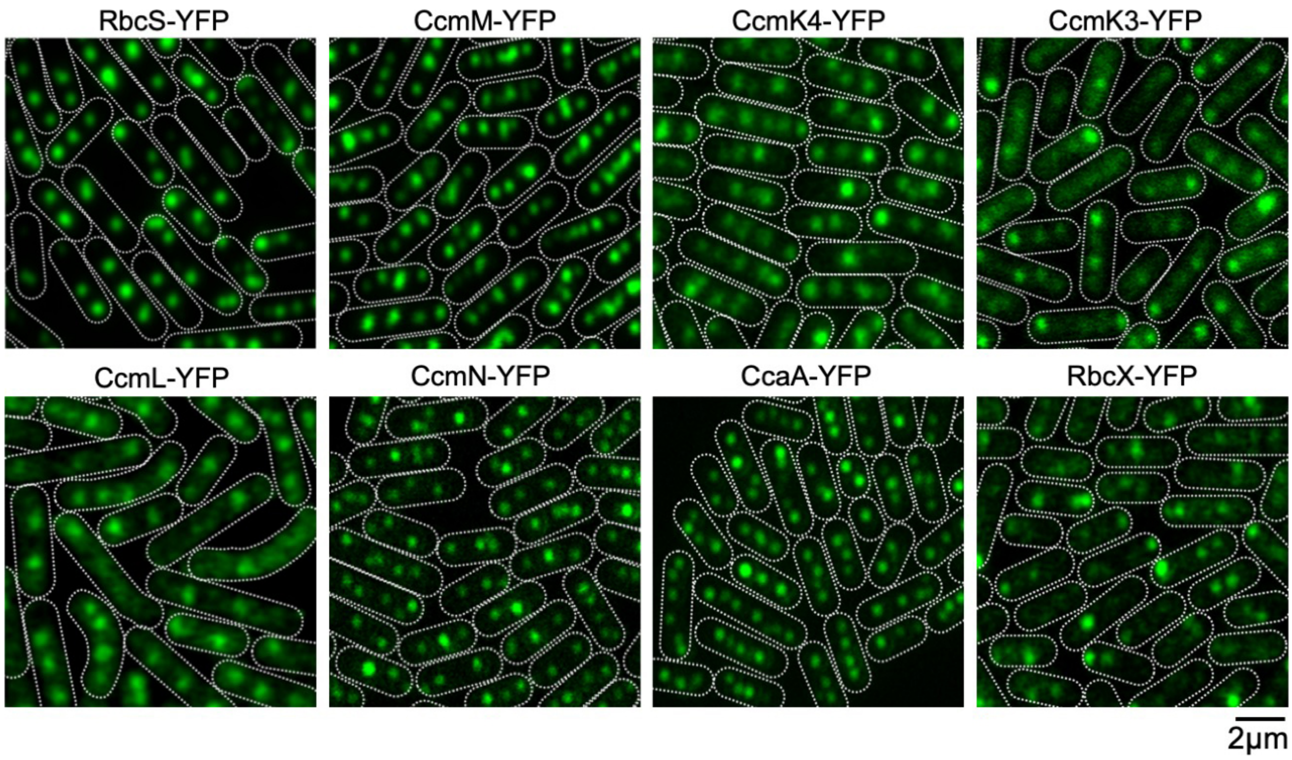


**Supplemental Figure 6. Confocal images of YFP-tagged cells.**
YFP signals are shown in green, indicative of YFP-labeled carboxysomes. White dash lines indicate cell borders.

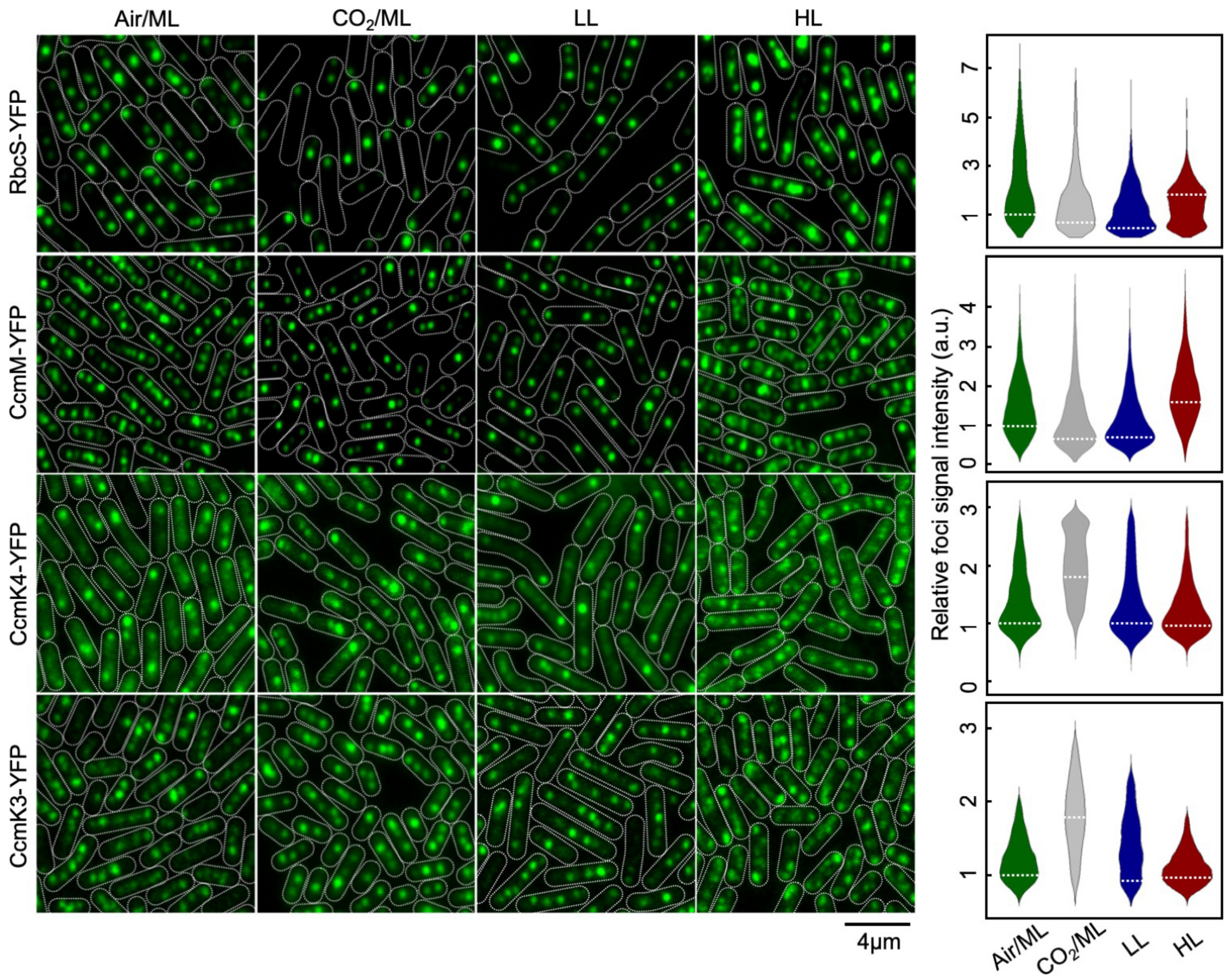


**Supplemental Figure 7. Confocal images of RbcS-YFP, CcmM-YFP, CcmK4-YFP and CcmK3-YFP cells under Air/ML, $CO_2$, LL, and HL and distribution profiles of carboxysome protein signal intensity.**

Imaging on the same strain under the four treatments was performed with the same parameters, allowing direct comparison of fluorescence intensities. Carboxysome and cell borders (shown as green and white dash lines, respectively) were optimized for presentation. The peak values obtained from kernel density fitting of the intensity distribution, as shown by white dashed lines, were used for examining changes in the abundance of the same carboxysome proteins under different conditions. Scale bar indicates 4 μm.

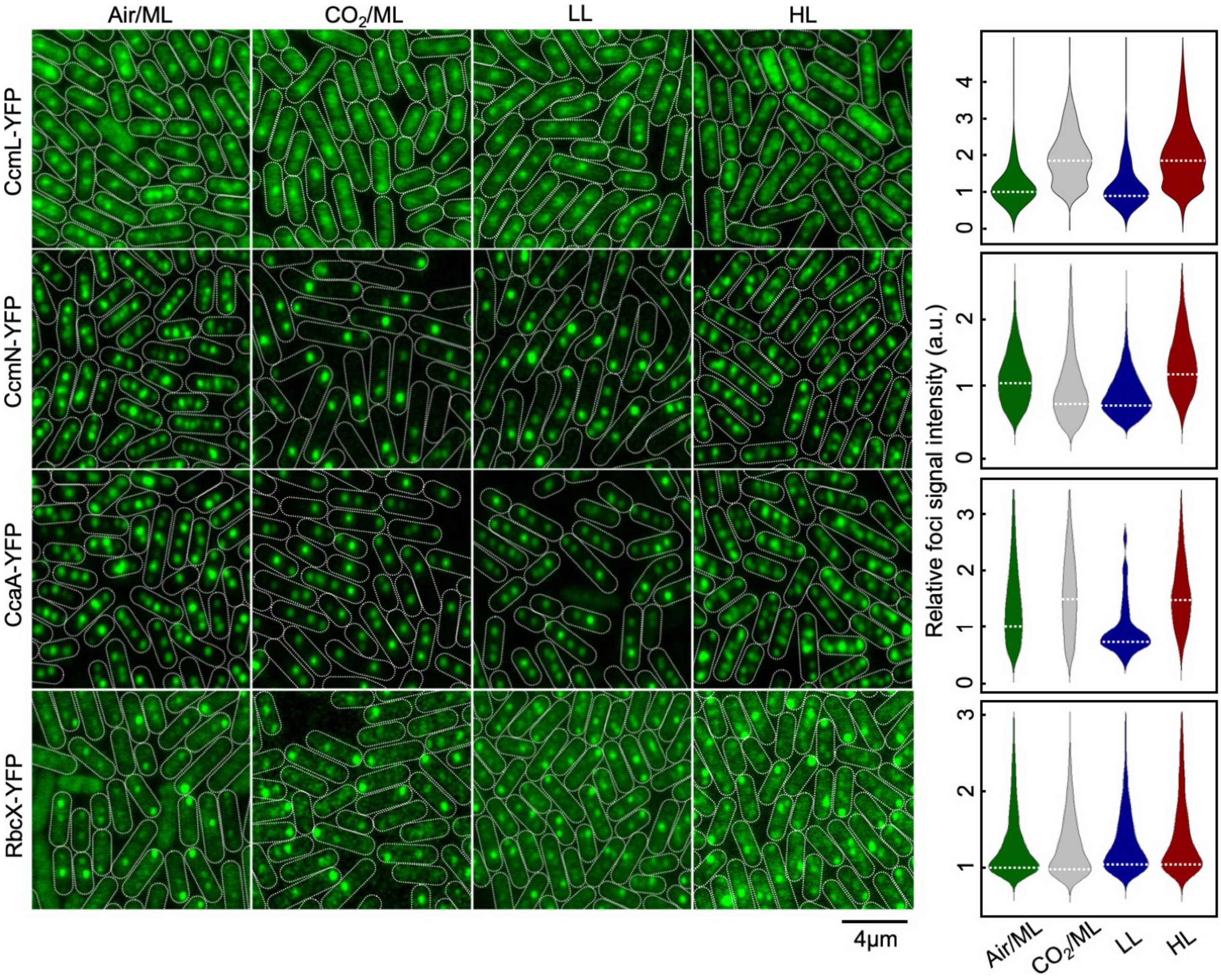


**Supplemental Figure 8. Confocal images of CcmL-YFP, CcmN-YFP, CcaA-YFP and RbcX-YFP cells under Air/ML, $CO_2$, LL, and HL and distribution profiles of carboxysome protein signal intensity (continuing Supplemental Figure 7).**

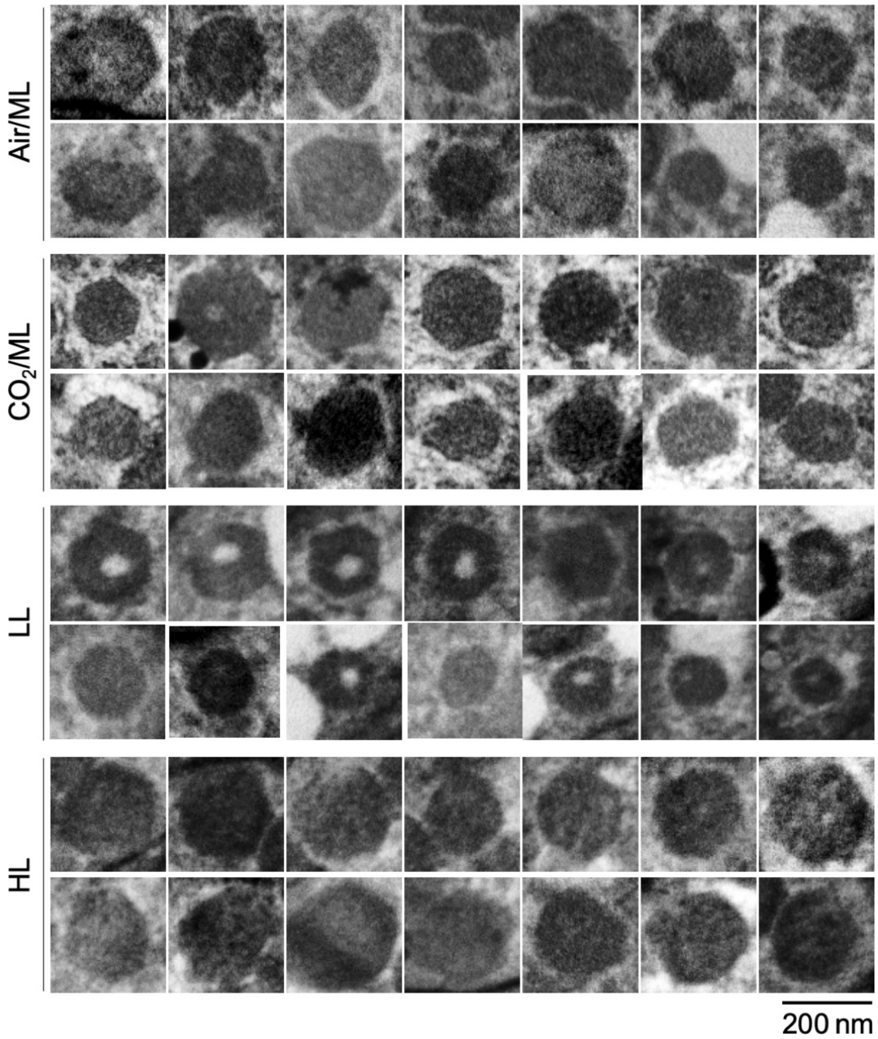


**Supplemental Figure 9. Thin-section EM images of WT Syn7942 cells under Air/ML, $CO_2$/ML, LL and HL.** The size of carboxysomes in cells grown under different environmental conditions varies. Low density regions were often found in the lumen of carboxysomes synthesized under LL (59%, 16 out of 27 β-carboxysomes), compared with 9% for Air/ML (3 out of 33), 12% for $CO_2$/ML (3 out of 25) and 8% for HL (4 out of 51).

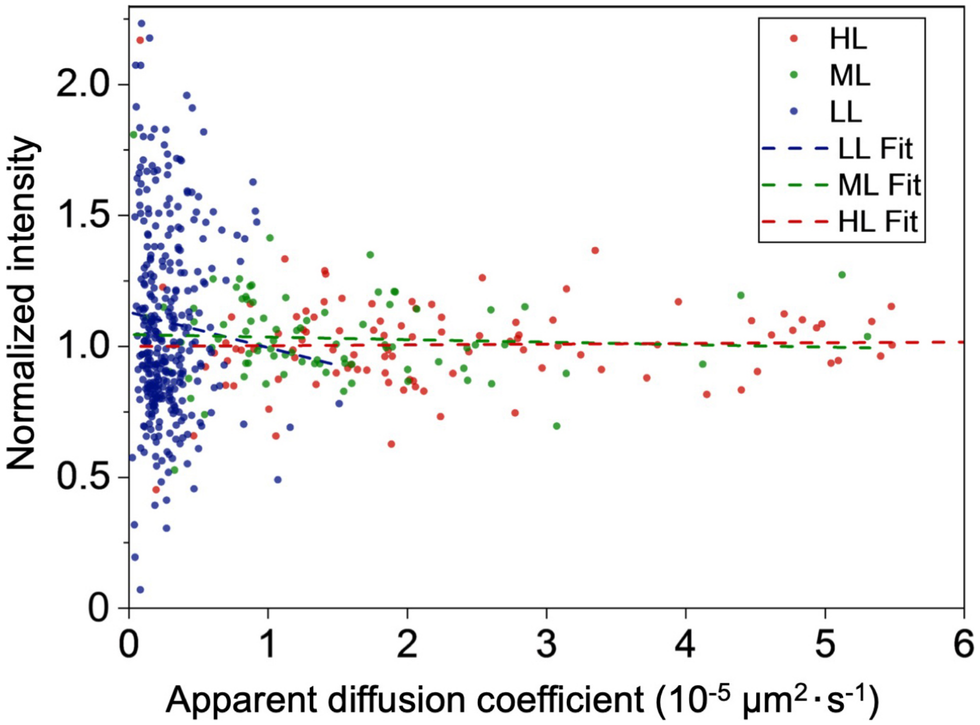


**Supplemental Figure 10. Changes in the diffusion coefficient of carboxysomes in Syn7942 cells under HL, ML and LL are not dependent on the carboxysome size*.***
No explicit correlation between diffusion coefficient and RbcL-YFP fluorescence intensities per carboxysome (representing the carboxysome size) under HL, ML and LL was observed (linear fitting $R^2 < 0.005$ for HL, ML and LL). 105, 84 and 336 carboxysomes were tracked under HL, ML and LL respectively.

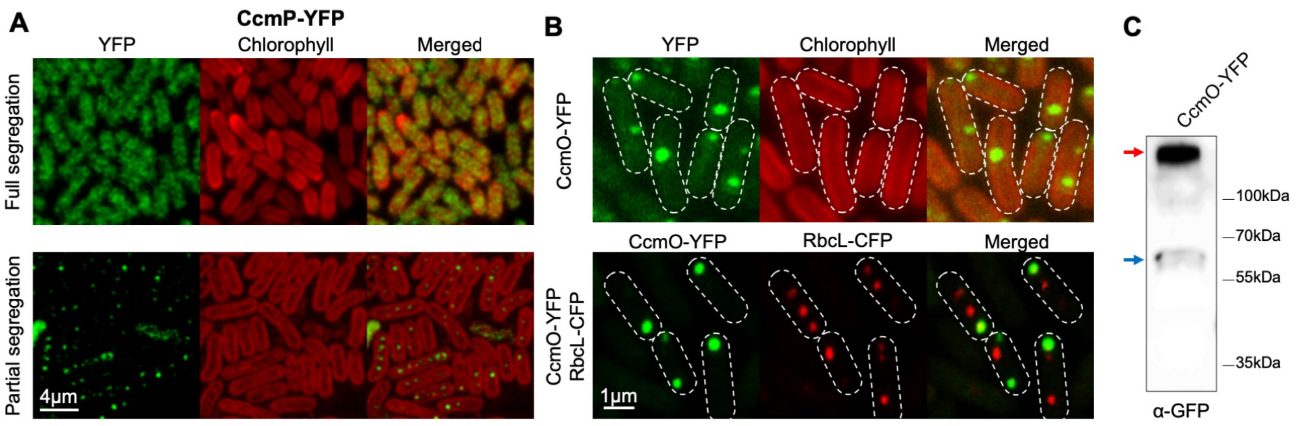


**Supplemental Figure 11. CcmP-YFP and CcmO-YFP Syn7942 cells.**

**(A)** Fully-segregated CcmP-YFP cells show abnormal carboxysome distribution and assembly, whereas partially-segregated cells present typical carboxysome distribution. Scale Bar indicates 4 µm.

**(B)** Only 1-2 fluorescence foci are seen in the fully-segregated CcmO-YFP cells fusion. Confocal imaging of the double-labeled CcmO-YFP::RbcL-CFP strain reveals that CcmO-YFP and RbcL-CFP are not strictly co-localized in the cell. Scale Bar indicates 1 µm.

**(C)** Immunoblot analysis of the soluble fractions of the CcmO-YFP strain cell lysates using an anti-fluorescent protein antibody, showing the presence of large aggregates of CcmO that failed to run into the gel, marked by the red arrow. The minor fractions of CcmO that run into the gel are marked by the blue arrow. For all western results, at least 3 independent experiments were conducted.

**Supplemental Table 1. Cell growth, carbon fixation and cell dimensions of Syn7942 WT and YFP-fusion mutants under Air/ML.**
Results are represented as mean ± SD. The numbers of cells in the cell-dimension analysis were shown as *n*. No significant differences in growth rate were observed, indicating that YFP tagging has no notable effects on cell growth and physiology.

| **Strains** | **Doubling time (hours) (*n* = 4)** | **Carbon fixation (nmol·ml$^{-1}$·min$^{-1}$) (*n* = 3)** | **Cell length (µm)** | **Cell width (µm)** | ***n*** |
|---|---|---|---|---|---|
| WT Syn7942 | 16.82 ± 1.31 | 3.91 ± 0.08 | 3.38 ± 1.65 | 0.88 ± 0.14 | 1360 |
| CcmK2-YFP | 17.26 ± 0.83 | 3.45 ± 0.09 | 3.4 ± 1.28 | 0.82 ± 0.1 | 951 |
| CcmK3-YFP | 18.03 ± 1.22 | 4.22 ± 1.06 | 3.66 ± 1.43 | 0.78 ± 0.08 | 1015 |
| CcmK4-YFP | 17.68 ± 1.44 | 3.78 ± 0.86 | 3.6 ± 1.45 | 0.82 ± 0.08 | 904 |
| CcmL-YFP | 18.59 ± 1.41 | 3.76 ± 0.47 | 3.16 ± 1.66 | 0.82 ± 0.17 | 668 |
| CcaA-YFP | 16.79 ± 0.98 | 3.38 ± 0.36 | 3.45 ± 1.39 | 0.81 ± 0.09 | 622 |
| CcmN-YFP | 17.27 ± 1.08 | 3.68 ± 0.68 | 3.34 ± 1.07 | 0.87 ± 0.07 | 751 |
| CcmM-YFP | 17.74 ± 1.02 | 3.76 ± 0.47 | 3.3 ± 1.53 | 0.86 ± 0.14 | 705 |
| RbcL-YFP | 17.78 ± 1.88 | 4.01 ± 0.62 | 3.22 ± 1.19 | 0.85 ± 0.1 | 556 |
| RbcS-YFP | 18.33 ± 2.13 | 3.70 ± 1.01 | 3.03 ± 1.07 | 0.78 ± 0.09 | 236 |
| RbcX-YFP | 18.14 ± 0.77 | 3.67 ± 0.87 | 3.67 ± 1.52 | 0.82 ± 0.12 | 799 |

**Supplemental Table 2. Immunoblotting estimation of the stoichiometry of carboxysomal proteins in cell lysates.**
The ratio of RbcL/S is 8:5.8 ± 0.8 ($n$ = 4), close to the previously proposed $RbcL_8S_5CcmM35$ assembly of Rubisco complexes in Syn7942 (Long et al., 2011). Quantification of CcmO and CcmN was not included, as they could form large aggregations in SDS-PAGE (Supplemental Figure 2C and Supplemental Figure 11C) and their contents could not be accurately estimated. Data are represented as mean ± SD from at least 3 biological repeats.

| **Category** | **Protein** | **Relative content (%)** |
|---|---|---|
| Shell proteins | CcmK2 | 70.4 ± 7.1 |
| | CcmK3 | 1.3 ± 0.1 |
| | CcmK4 | 2.0 ± 0.2 |
| | CcmL | 0.2 ± 0.01 |
| Structural proteins | CcmM | 2.5 ± 0.5 |
| CA enzymes | CcaA | 1.7 ± 0.4 |
| Rubisco enzymes | RbcL | 12.6 ± 3.5 |
| | RbcS | 9.1 ± 0.8 |
| Rubisco chaperone | RbcX | 0.1 ± 0.01 |

**Supplemental Table 3. Carboxysome content per cell under Air/ML, $CO_2$/ML, LL and HL determined by confocal imaging.**

| Strains | Air/ML | $CO_2$/ML | LL | HL |
|---|---|---|---|---|
| CcmK3-YFP | 3.49 ± 1.94 (*n* = 538) | 2.18 ± 2.33 (*n* = 648) | 2.63 ± 1.23 (*n* = 542) | 5.82 ± 3.15 (*n* = 295) |
| CcmK4-YFP | 3.12 ± 1.52 (*n* = 410) | 1.73 ± 1.86 (*n* = 397) | 2.15 ± 0.93 (*n* = 514) | 5.65 ± 2.76 (*n* = 303) |
| CcmL-YFP | 3.40 ± 1.89 (*n* = 326) | 1.55 ± 1.71 (*n* = 574) | 1.98 ± 0.96 (*n* = 587) | 7.53 ± 4.88 (*n* = 720) |
| CcmM-YFP | 4.77 ± 2.51 (*n* = 565) | 1.79 ± 1.92 (*n* = 599) | 2.12 ± 1.04 (*n* = 354) | 7.88 ± 2.57 (*n* = 565) |
| CcmN-YFP | 4.35 ± 2.71 (*n* = 572) | 1.61 ± 1.73 (*n* = 579) | 2.19 ± 1.16 (*n* = 566) | 6.65 ± 3.28 (*n* = 424) |
| RbcS-YFP | 3.43 ± 1.8 (*n* = 359) | 1.22 ± 1.31 (*n* = 343) | 2.25 ± 1.49 (*n* = 130) | 5.45 ± 2.71 (*n* = 385) |
| RbcX-YFP | 2.58 ± 1.11 (*n* = 552) | 2.02 ± 2.16 (*n* = 700) | 2.48 ± 1.03 (*n* = 104) | 4.75 ± 2.66 (*n* = 366) |
| CcaA-YFP | 3.18 ± 1.23 (*n* = 334) | 2.11 ± 2.33 (*n* = 361) | 2.90 ± 1.50 (*n* = 422) | 6.45 ± 3.95 (*n* = 427) |

**Supplemental Table 4. Evaluation and quality control of quantitative microscopy.**
To ensure adequate sampling for accurate estimation of peak carboxysomal signal intensities from highly heterogeneous populations, evaluation was performed for data shown in Figures 2 and 3, Table 1 and Supplemental Figure 7-8. Observation values (as peak intensities of individual carboxysomal signals that scaled to the gold-standard copy number reference to data from Slimfield Air/ML) were randomly distributed into at least 3 groups that contain no less than 70 entries each. The sampling errors were determined from the standard error (SE) of mean group peak intensities. The maximum errors as SE/Peak intensity were controlled below 5%.

| Protein | Treatment | Mean group peak value | SE of Peak value within groups | Number of randomized groups | Group sample counts | Normalized SE (SE/Peak, %) |
|---|---|---|---|---|---|---|
| CcmK3 | Air/ML | 92 | 0.7 | 23 | 100 | 0.80 |
| | $CO_2$/ML | 172 | 1.5 | 20 | 100 | 0.88 |
| | LL | 83 | 1.0 | 15 | 100 | 1.16 |
| | HL | 87 | 0.6 | 21 | 100 | 0.65 |
| CcmK4 | Air/ML | 314 | 3.4 | 17 | 100 | 1.07 |
| | $CO_2$/ML | 562 | 5.7 | 19 | 100 | 1.02 |
| | LL | 313 | 4.9 | 17 | 100 | 1.57 |
| | HL | 304 | 2.3 | 32 | 100 | 0.76 |
| CcmL | Air/ML | 37 | 1.5 | 3 | 100 | 4.05 |
| | $CO_2$/ML | 66 | 1.2 | 3 | 100 | 1.81 |
| | LL | 34 | 0.7 | 3 | 100 | 2.05 |
| | HL | 69 | 1.8 | 3 | 70 | 2.61 |
| CcmM | Air/ML | 719 | 8.9 | 36 | 100 | 1.23 |
| | $CO_2$/ML | 468 | 8.8 | 23 | 100 | 1.87 |
| | LL | 483 | 7.6 | 36 | 100 | 1.57 |
| | HL | 1176 | 12.8 | 23 | 100 | 1.09 |
| CcmN | Air/ML | 74 | 0.7 | 26 | 100 | 0.94 |
| | CO2/ML | 52 | 0.4 | 31 | 100 | 0.78 |
| | LL | 51 | 0.3 | 40 | 100 | 0.61 |
| | HL | 82 | 0.6 | 50 | 100 | 0.72 |
| CcaA | Air/ML | 86 | 0.9 | 33 | 100 | 1.07 |
| | $CO_2$/ML | 129 | 2.0 | 13 | 100 | 1.55 |
| | LL | 65 | 2.9 | 3 | 70 | 4.40 |
| | HL | 122 | 1.1 | 28 | 100 | 0.91 |
| RbcS | Air/ML | 6822 | 125.9 | 15 | 100 | 1.85 |
| | $CO_2$/ML | 4401 | 136.5 | 8 | 100 | 3.10 |
| | LL | 2934 | 111.5 | 7 | 100 | 3.80 |
| | HL | 12057 | 232.0 | 19 | 100 | 1.92 |
| RbcX | Air/ML | 39 | 0.2 | 13 | 100 | 0.58 |
| | $CO_2$/ML | 38 | 0.4 | 13 | 100 | 0.96 |
| | LL | 40 | 0.5 | 14 | 100 | 1.28 |
| | HL | 40 | 0.3 | 18 | 100 | 0.87 |

**Supplemental Table 5. Carboxysome properties in Syn7942 varies under Air/ML, $CO_2$/ML, LL and HL, determined by Slimfield, confocal and EM imaging.**
*Data estimated using calculations suggested in the previous model (Long et al., 2011).

| Properties of average carboxysome | Air/ML | $CO_2$/ML | LL | HL |
|---|---|---|---|---|
| Diameter (nm) | 192 | 144 | 151 | 208 |
| Surface area ($10^4$ $nm^2$)* | 8.82 | 4.96 | 5.46 | 10.36 |
| Internal volume ($10^6$ $nm^3$)* | 1.88 | 0.75 | 0.87 | 2.42 |
| Ratio of surface area:volume* | 0.039 | 0.052 | 0.050 | 0.036 |
| Copies of Rubisco<br>(Based on RbcS content) | 853 | 550 | 367 | 1507 |
| Rubisco density in carboxysome lumen<br>($10^{-4}$ copy/$nm^3$) | 4.54 | 7.33 | 4.21 | 6.23 |
| Ratio of CcmM:Rubisco | 0.843 | 0.851 | 1.316 | 0.780 |
| Ratio of CcaA:Rubisco | 0.101 | 0.235 | 0.177 | 0.081 |
| Ratio of CcmK4:CcmK3 | 3.8 | 3.6 | 4.1 | 3.9 |
| Coverage of CcmK3-K4 +CcmL on surface (%)* | 19.2 | 34.1 | 31.0 | 16.3 |
| Carbon fixation rate per Rubisco (a.u.) | 1.00 | 7.15 | 1.80 | 0.83 |

**Supplemental Table 6. Accession numbers for genes in this work**

| Gene | Locus tag | Gene ID for corresponding protein |
|---|---|---|
| *ccmK3* | *synpcc7942_0284* | ABB56316 |
| *ccmK4* | *synpcc7942_0285* | ABB56317 |
| *ccmP* | *synpcc7942_0520* | ABB56552 |
| *ccmK2* | *synpcc7942_1421* | ABB57451 |
| *ccmL* | *synpcc7942_1422* | ABB57452 |
| *ccmM* | *synpcc7942_1423* | ABB57453 |
| *ccmN* | *synpcc7942_1424* | ABB57454 |
| *ccmO* | *synpcc7942_1425* | ABB57455 |
| *rbcL* | *synpcc7942_1426* | ABB57456 |
| *rbcS* | *synpcc7942_1427* | ABB57457 |
| *ccaA* | *synpcc7942_1447* | ABB57477 |
| *rbcX* | *synpcc7942_1535* | ABB57565 |

**Supplemental Table 7. PCR primers used in this study for gene cloning and sequencing.**

| Primers | Sequence (5'-3') |
|---|---|
| CcmK3_F | TCAAGCTGTTGGGCGAAGTC |
| CcmK3_R | AAGCATTGGCTTCGGCATCC |
| CcmK4_F | AGCGGTGATTGCAAGACACG |
| CcmK4_R | AACGCCAAAGCCACCATGAG |
| CcmK2_F | GCGGGAAGCTCAAGTTTGGG |
| CcmK2_R | ACTCGCTGGCCTATCCAGAC |
| CcmL_F | GAGCGAAGCTGTTACCTCAC |
| CcmL_R | GTTCCGTCCCAATCCGATAG |
| CcmM_F | ACGTGGACTCATGCCTATTG |
| CcmM_R | GGCAGAATAGTGCTGGAAAC |
| CcmN_F | CGAACACGAAGGTGAATACG |
| CcmN_R | AGTTCCGGTAGCTCCAACAG |
| CcmO_F | CGATGTCGCTGGACAGGTTC |
| CcmO_R | GTCGCGCTCGACTTGGATAC |
| CcmP_F | CTGTTTGCCACCGATGACAG |
| CcmP_R | CGCTCATGTTCAGCCAACTC |
| RbcL_F | CAACCTCGAAGCCGTGTTGC |
| RbcL_R | GGCTGTTAGCTCCGACTCTG |
| RbcS_F | ACGGGATTCACTTCCGTGTC |
| RbcS_R | GTGTTCCCTTGCCGTCGTAG |
| CcaA_F | AAACAGAGCTCGCCGTTATC |
| CcaA_R | GCATGAACGGGCATTCATTG |
| RbcX_F | AAGCAGGTTGGCAGCCTATC |
| RbcX_R | TCGCTGTCATCAAGGCATCG |
| CcmK3_YFP_F | CCGATCGACTTCACGGCTGAATCCGAGCCCTTTCGGTCTCTGCCG GGCCCGGAGCTGCC |
| CcmK3_YFP_R | ACTCCTTCGCGTCGCAGTGTCGATTGTAAAGTCCGTGGCATTCCG GGGATCCGTCGACC |
| CcmK4_YFP_F | TATCGACTCTCTGCCGAAGGAACTGGTAGTGGCCGCCGTCTGCCG GGCCCGGAGCTGCC |
| CcmK4_YFP_R | CCCTCAGCCCAAATCAACCCTTTTAATCAGTCGCTACCCATTCCGG GGATCCGTCGACC |
| CcmK2_YFP_F | CCGATTCGCTACACCGAAGCTGTTGAACAATTCCGCATGCTGCCG GGCCCGGAGCTGCC |
| CcmK2_YFP_R | GATGAAAGGGCTATTGAAACGACTCAACAGCCCTCAACCATTCCG GGGATCCGTCGACC |
| CcmL_YFP_F | GTGGAAAACCGCTCCGTCTACGACAAACGCGAGCACAGCCTGCC GGGCCCGGAGCTGCC |
| CcmL_YFP_R | ATCCAGATCAATGACCAGCAGGGATTCAATCCCTGCCCAATTCCG GGGATCCGTCGACC |
| CcmM_YFP_F | GCTCGTCGTCGCGTGGCTGAACTGTTGATTCAAAAGCCGCTGCCG GGCCCGGAGCTGCC |
| CcmM_YFP_R | TGCATGAAGACAAGGGCGATCGCCGGATTCCCGGCACAGATTCC GGGGATCCGTCGACC |
| CcmN_YFP_F | CAGTTTTTGCGGATGCGCCAGAGCATGTTCCCCGATCGCCTGCCG GGCCCGGAGCTGCC |
| CcmN_YFP_R | ACAAAGTACCCTGCAGCTCCTAGAGCTGCTGTGCACATCATTCCG GGGATCCGTCGACC |
| CcmO_YFP_F | GCCGAGCCCTTAGAGCTCCCCAATCCTCGTGATGATCAGCTGCCG GGCCCGGAGCTGCC |
| CcmO_YFP_R | TTTCAGCCAAAAACCAAGGGAATTACAGTGCTTTTTCCAATTCCGG GGATCCGTCGACC |

| | |
|---|---|
| CcmP_YFP_F | CTGTCGGGACGTGAGCATCCCGGCGATCGCTCGCGGGAGCTGCC GGGCCCGGAGCTGCC |
| CcmP_YFP_R | GCCGCAGAAATGCCAGATGTTCTGCATTCGCCATGACCGATTCCG GGGATCCGTCGACC |
| RbcL_YFP_F | AAAGAGATCAAGTTCGAATTCGAAACGATGGACAAGCTCCTGCCG GGCCCGGAGCTGCC |
| RbcL_YFP_R | GCAGCAACGCTCACTCCCCCAGCGATAGTCAGAGGCTCCATTCCG GGGATCCGTCGACC |
| RbcS_YFP_F | CAAACCGTGAGCTTCATCGTTCATCGTCCCGGCCGCTACCTGCCG GGCCCGGAGCTGCC |
| RbcS_YFP_R | TGCCCAAAGGCAGGAGCAGCTATCAAGACAAATCAGGCTATTCCG GGGATCCGTCGACC |
| CcaA_YFP_F | CCTGAACAACAACAGCGGATTTATCGCGGCAATGCTAGCCTGCCG GGCCCGGAGCTGCC |
| CcaA_YFP_R | ACACAGGCAGATCTCAGCAGGGTCGAAGATGCTTCGATCATTCCG GGGATCCGTCGACC |
| RbcX_YFP_F | GCCACTCCTGATGATGCTTCTAATGCCTCCCATGCGGATCTGCCG GGCCCGGAGCTGCC |
| RbcX_YFP_R | ATCGCGGCAGGCCCTTCAAAATCAACGTGTTGAACAATTATTCCGG GGATCCGTCGACC |
| CcmK3_insert_segF | ATCCCGAATCCACAAGAA |
| CcmK3_insert_segR | GCCCGCATGTAGCTCACAAT |
| CcmK4_insert_segF | GCCGAAGGAACTGGTAGTGG |
| CcmK4_insert_segR | GGGCTTCATCCCAGACATAG |
| CcmK2_insert_segF | GTTCTCCCGATTCGCTACA |
| CcmK2_insert_segR | AACAAATCTGCCAAGGTGC |
| CcmL_insert_segF | AAACCGCTCCGTCTACGAC |
| CcmL_insert_segR | TTGTTGGGCTCGGCATTCT |
| CcmM_insert_segF | GGATCGAGTTCGAGCGATGT |
| CcmM_insert_segR | TCACCACTGGCAAAGTAGCG |
| CcmN_insert_segF | CTAAGGTCTACGGCAAGGAA |
| CcmN_insert_segR | GCTGAGAATAGGCGGGAAG |
| CcmO_insert_segF | TGAGCCGCAGTCGGATAG |
| CcmO_insert_segR | GAAGGCGTTGACATCACTCT |
| CcmP_insert_segF | TCAGTGCGATTGGCAGTTT |
| CcmP_insert_segR | TCATTCCCGATCTTCGCCA |
| RbcL_insert_segF | CGTGAAGCTGGCAAGTGG |
| RbcL_insert_segR | GGAGGCAGGTACGAGAAAGT |
| RbcS_insert_segF | CAACATCAAGCAGTGCCAAAC |
| RbcS_insert_segR | CACCACCAATCACTCCAACG |
| CcaA_insert_segF | CCCTGACCGAAGAAGTGGC |
| CcaA_insert_segR | CAGGTAAGTCGGCAGGAAG |
| RbcX_insert_segF | GCCACTCCTGATGATGCTTC |
| RbcX_insert_segR | CACTTTGGGCGAGGGAGATT |
